# “Target-and-release” nanoparticles for effective immunotherapy of metastatic ovarian cancer

**DOI:** 10.1101/2024.07.05.602135

**Authors:** Ivan S. Pires, Gil Covarrubias, Victoria F. Gomerdinger, Coralie Backlund, Apoorv Shanker, Ezra Gordon, Shengwei Wu, Andrew J. Pickering, Mariane B. Melo, Heikyung Suh, Darrell J. Irvine, Paula T. Hammond

## Abstract

Immunotherapies such as checkpoint inhibitors (CPI) are effective in treating several advanced cancers, but these treatments have had limited success in metastatic ovarian cancer (OC). Here, we engineered liposomal nanoparticles (NPs) carrying a layer-by-layer (LbL) polymer coating that promotes their binding to the surface of OC cells. Covalent anchoring of the potent immunostimulatory cytokine interleukin-12 (IL-12) to phospholipid headgroups of the liposome core enabled the LbL particles to concentrate IL-12 in disseminated OC tumors following intraperitoneal administration. Shedding of the LbL coating and serum protein-mediated extraction of IL-12-conjugated lipids from the liposomal core over time enabled IL-12 to disseminate in the tumor bed following rapid NP localization in tumor nodules. Optimized IL-12 LbL-NPs promoted robust T cell accumulation in ascites and tumors in mouse models, extending survival compared to free IL-12 and remarkedly sensitizing tumors to CPI, leading to curative treatments and immune memory.

## Introduction

Ovarian cancer (OC) treatment is particularly challenging due to late diagnosis and metastatic spread.^1,2^ A promising approach for late-stage cancer treatment is immunotherapy.^3–5^ However, poor baseline lymphocyte infiltration and an immunosuppressive tumor microenvironment (TME) have correlated with limited benefits of immunotherapy in OC patients.^6,7^ Immunostimulatory agents such as cytokines and costimulatory antibodies may have the potential to overcome these limitations, but systemic administration of these therapeutics is severely limited by dose-limiting toxicities.^8,9^

Nanoparticles (NPs) are promising vehicles to deliver immunotherapeutics.^10,11^ In OC, NPs have been used to deliver various immunomodulatory agents such as nucleic acids^12,13^, proteins^14^, and small molecules^15,16^. We previously reported that NPs coated with poly-L-arginine (PLR)/poly-L-glutamate (PLE) bilayers via layer-by-layer (LbL) deposition showed selective binding to the surface of OC cells.^17^ In mouse OC models, administration of liposomal LbL-NPs carrying the potent immunostimulatory cytokine interleukin-12 (IL-12) showed reduced toxicity over systemic IL-12 dosing but only modest therapeutic efficacy.^18,19^ We hypothesized that the non-covalent nickel-histidine interaction used to tether IL-12 to these particles was very short-lived *in vivo*,^20,21^ leading to premature release of the cytokine before uptake in tumors.

Here, we demonstrate that covalent conjugation of IL-12 to the liposomal core of LbL-NPs greatly improves targeting and retention of IL-12 in peritoneally-disseminated OC tumors, enabling immunological and therapeutic effects not observed with free cytokine treatment. Mechanistic investigations revealed that these LbL-NPs rapidly accumulated in tumor nodules upon intraperitoneal (i.p.) administration, followed by shedding of the LbL coating and gradual release of IL-12-lipid conjugates via lipid extraction by serum proteins present in interstitial fluid. Both rapid LbL-mediated cancer cell targeting and slow cytokine release with sustained retention on cell membrane surfaces were crucial for therapeutic efficacy. These findings demonstrate the potential of “target and release” NP designs to effectively concentrate cytokine in disseminated ovarian cancer lesions and promote robust anti-tumor immunity.

## Results and Discussion

### Dynamics of IL-12-conjugated LbL-NPs on contact with physiologic fluids

The overall design of the LbL-NP system is shown in **Fig. 1a**. An immunostimulatory payload (here, a single-chain version of the potent cytokine IL-12) is linked to the surface of a liposomal core particle, followed by sequential LbL deposition of a layer of positively charged poly-L-arginine (PLR) and then a layer of negatively-charged poly-L-glutamate (PLE). To understand how the stability of the IL-12/NP association impacts the efficacy of this system, we compared the previously employed non-covalent nickel:polyhistidine (Ni) interaction^18^ to a new formulation with a N-aryl maleimide (Mal)-cysteine linkage of IL-12 to the particles which forms a stable covalent bond^22^ (**Supplementary Table 1, Fig. S1a-c**). IL-12-conjugated NPs were synthesized with both linker chemistries (Ni or Mal) in either unlayered (UL) or PLR/PLE-layered (LbL) formats. Ni and Mal NPs had similar sizes (**Fig. S2a**), zeta potentials (**Fig. S2b**), yields (>70%), and loadings of IL-12 (∼10-13 wt%, corresponding to ∼50 IL-12 molecules/particle^18^ , **Fig. S2c-e**).

**Figure 1.**
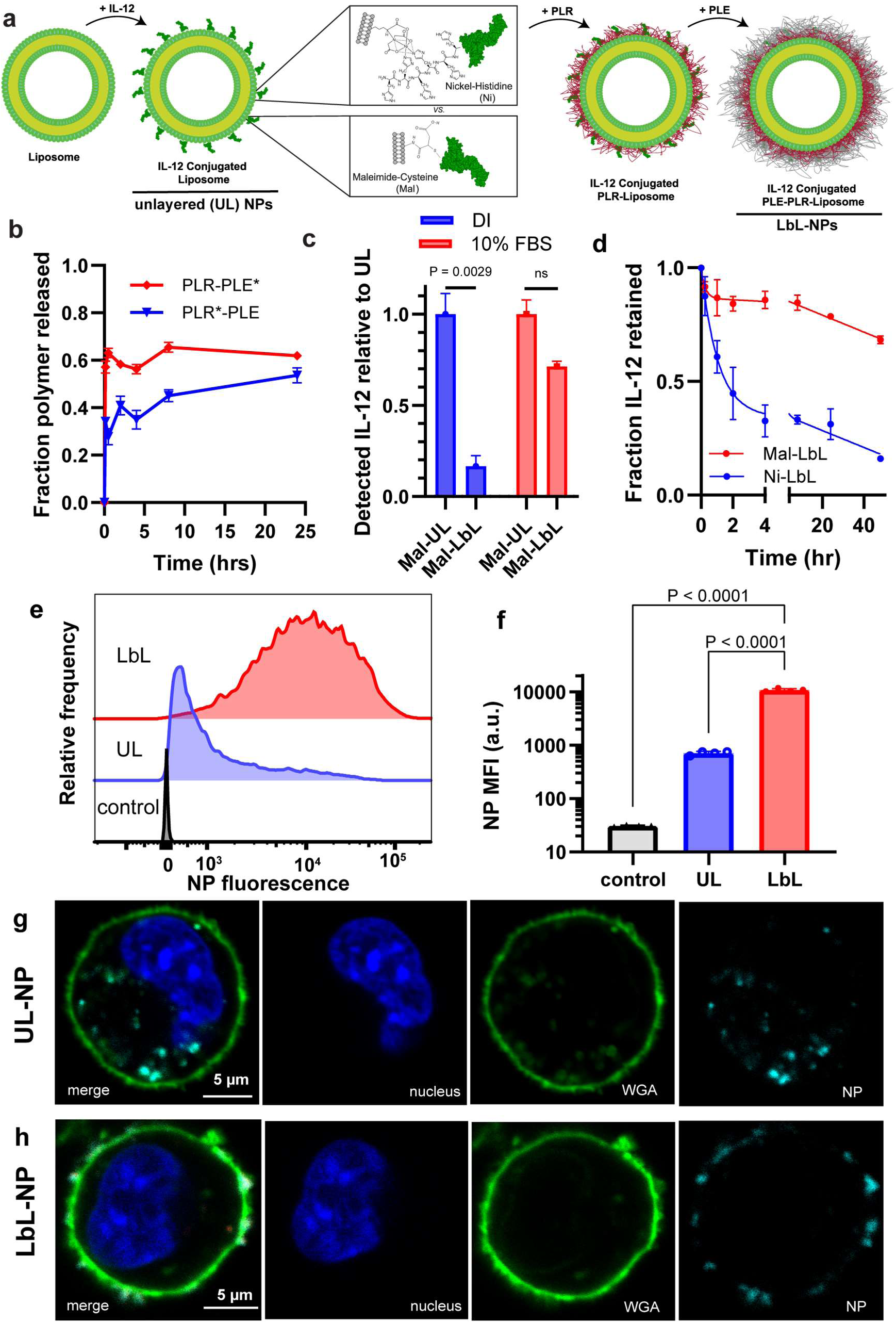
LbL NPs undergo dynamic reorganization on contact with physiologic fluids. **a,** Schematic for assembly of LbL-NPs with either Mal or Ni linker chemistries for conjugation of IL-12 onto NPs. **b,** Quantification of total PLE and PLR retained with LbL-NPs upon incubation in cell-free ascites fluid at 37 °C (* indicates fluorophore tagged polymer) (mean ± s.e.m.). **c,** Quantification of total IL-12 available for monoclonal antibody binding from Mal NPs either in diH2O or 10% FBS media (mean ± s.e.m.). **d,** Quantification of total IL-12 released from LbL-NPs upon incubation in cell-free ascites fluid at 37 °C (mean ± s.e.m.). **e,** Representative flow cytometry fluorescence histogram of HM-1 cells incubated with unlayered (UL) or PLE-coated LbL-NPs for 4 hours *in vitro*. **f,** Quantification of median fluorescence intensity (MFI) of treated HM-1 cells from e (mean ± s.d.). **g-h,** Representative confocal images of HM-1 cells incubated with UL (**g**) or LbL NPs (**h**) for 4 hours – UL images adjusted relative to LbL to visualize internalized NPs (blue, Hoechst 33342 nuclear stain; green , wheat germ agglutinin (WGA) cell surface stain; cyan, nanoparticles). Data are representative of at least two independent experiments. Statistical comparisons in **c** and **f** were performed using two and one-way analysis of variance (ANOVA), respectively, with Tukey’s multiple-comparisons test.

LbL-NPs of this type are stored in deionized water (diH2O) to maintain colloidal stability and are diluted in 5% dextrose for isosmotic *in vivo* administration.^17,19,23^ However, we expect rearrangements in the LbL film to occur *in vivo* due to the high ionic strength of the intraperitoneal fluid. To simulate the exposure of NPs to the tumor-bearing peritoneal space, we incubated LbL-NPs in cell-free ascites fluid collected from mice bearing tumors formed from the highly metastatic OC cell line OV2944-HM1 (HM-1).^24^ Using fluorescently-labeled polymers to assess the release of the LbL components, we found both PLR and PLE exhibited a burst release in the presence of ascites fluid from the particles of ∼40% and ∼60%, respectively; following this initial rearrangement the remaining LbL film was stable for at least 24 hr (**Fig. 1b**). Similar PLE release profiles were observed when particles were incubated in HEPES-buffered saline solution with or without serum supplementation (**Fig. S3a**), suggesting that this partial polymer shedding was driven by the change in ionic strength from assembly and storage in diH2O.

Given this reorganization of the LbL film on contact with physiological fluids, we next evaluated the accessibility of particle-bound IL-12 by capturing Mal NPs on microtiter plates and testing binding of an IL-12-specific monoclonal antibody via an enzyme-linked immunosorbent assay (ELISA, **Fig. S3b**). LbL coating of IL-12-conjugated liposomes reduced the accessibility of IL-12 conjugated to the as-synthesized particles in diH2O by ∼90%, but on exposure to serum-containing buffer, the majority of particle-bound cytokine was accessible to the anti-IL-12 antibody (**Fig. 1c**). Further, addition of either UL or LbL NPs to IL-12 reporter cells in the presence of serum elicited roughly equivalent IL-12 signaling (**Fig. S3c-d**). Exposure of NP-bound IL-12 enables IL-12 signaling and also opens the potential for release of the IL-12 from the particle core over time, either due to disruption of the Ni/his-tag interaction (for Ni particles) or via protein-mediated extraction of lipid-anchored IL-12 from the core liposomal bilayer (for Mal particles). We incubated LbL-NPs carrying fluorescently tagged IL-12 with ascites fluid collected from HM-1 tumor-bearing mice, and IL-12 release from the particles was measured over time. While Ni-LbL particles released more than 50% of their IL-12 payload in 2 hr, cytokine release from Mal-LbL particles was much slower, with ∼70% of the payload still bound to the particles at 48 hr (**Fig. 1d**). Finally, we assessed whether the LbL-NPs retained effective cancer cell targeting on exposure to physiologic conditions. When incubated with HM-1 cells in the presence of complete cell culture media, LbL-coated NPs showed a >10-fold increased association with the cancer cells relative to uncoated particles (**Fig. 1e-f**). We previously showed that LbL-NPs selectively bind to OC cells over healthy cells.^17,25,26^ Confocal imaging confirmed that Mal-LbL NPs were primarily located on the cell membrane 4 hours after NP dosing, while Mal-UL NPs were internalized in the same time frame, consistent with our prior studies of NPs carrying Ni-anchored IL-12 and suggesting that despite the initial partial shedding of polymer from the layered particles, the remaining LbL film still effectively mediated cancer cell surface targeting (**Fig. 1g-h**).

Altogether, these analyses predict a dynamic process of rapid LbL coating reorganization on injection *in vivo*, enabling IL-12 exposure that could promote immune engagement while retaining effective OC cell surface targeting. Linkage of the cytokine to particles via Ni:histag interactions will simultaneously lead to rapid release of IL-12 from the particle carrier, while covalently anchored IL-12 is predicted to exhibit a much slower release of lipid-bound IL-12.

### LbL-NPs rapidly associate with tumor tissue in vivo and covalent IL-12 conjugation enables prolonged cytokine retention in tumors

We next characterized the *in vivo* pharmacokinetics of IL-12 delivery via LbL-NPs in a model of disseminated OC.^24^ In this model, 7 days after i.p. inoculation of luciferase-expressing HM-1 cells (HM-1-luc), tumor nodules are detected across the intestines, omentum, and urogenital tract (UGT), with a preference for the omentum and UGT as is clinically observed with metastatic OC (**Fig. S4a-b**).^27^ Employing fluorescently tagged IL-12 and fluorescently labeled lipids in the NP formulations, we tracked the kinetics of NP and IL-12 clearance from the i.p. space using whole-animal fluorescence imaging. UL-NPs exhibited an exponential decay in NP signal over time, but LbL-NPs, by contrast, showed a partial clearance over 24 hr, followed by prolonged retention of ∼10% of particles from 24 hrs to 4 days (**Fig. 2a**). The kinetics of IL-12 clearance were affected both by the presence of the LbL coating and chemistry of NP linkage: IL-12 loaded on Ni-UL NPs cleared with kinetics identical to free IL-12 injected i.p., suggesting rapid release of the cytokine from the Ni-UL carrier (**Fig 2b)**. LbL coating of Ni NPs prolonged IL-12 persistence, but interestingly Mal-UL NPs provided a similar IL-12 retention in the i.p. space (**Fig. 2b-c**). Mal-LbL NPs exhibited the slowest IL-12 clearance, with ∼50% of the cytokine signal still present at 4 days post injection (**Fig. 2c**). Notably, IL-12 administered as Mal-LbL NPs had slower clearance than its lipid carrier, suggesting dissociation of the cytokine payload from the carrier particles over time, consistent with our *in vitro* analyses of NP stability.

**Figure 2.**
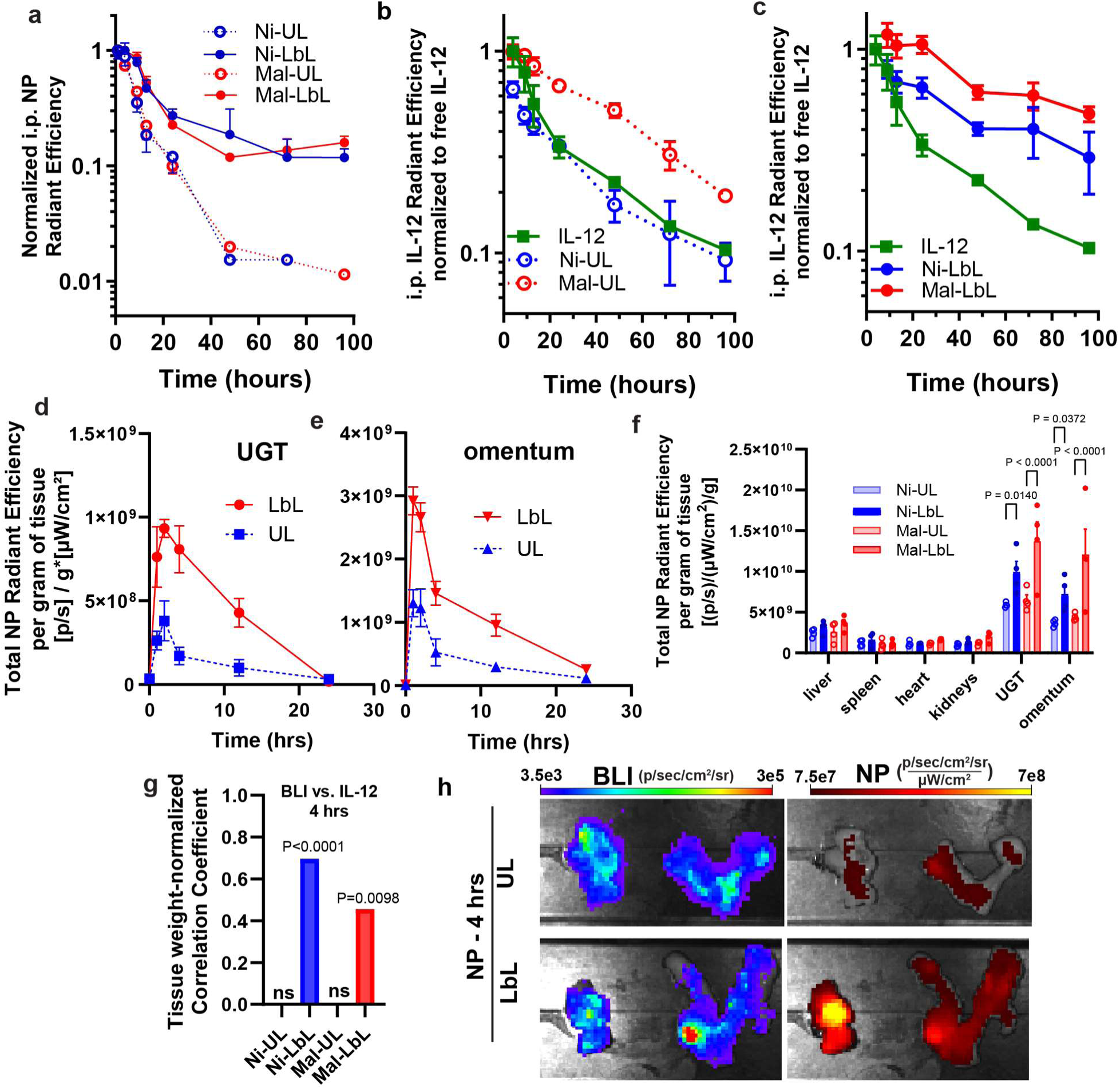
LbL coating enables targeting of tumor tissue *in vivo* and enhanced i.p. retention of NPs and IL-12. **a-c,** B6C3F1 mice (*n*=8/group for 0-24 hrs and n=3/group for 24-96 hrs) inoculated with 10^6^ HM-1-luc tumor cells on day 0 were administered fluorescently-tagged NPs carrying 20 µg IL-12 (or an equivalent dose of free IL-12) on day 14. Shown are whole-animal imaging NP fluorescence **(a)** and IL-12 fluorescence **(b, c)** from the i.p. space collected over time post dosing (mean ± s.e.m.). **d-e,** B6C3F1 mice (n=4/group) inoculated with 10^6^ HM-1-luc tumor cells on day 0 were administered 100 µg fluorescently-tagged LbL-NPs or UL-NPs (devoid of IL-12) on day 14. UGT and omentum tissue were harvested at 1, 2, 4, 12 and 24 hrs after dosing and imaged ex-vivo via IVIS. Shown are weight-normalized tissue NP fluorescence in **d** UGT and **e** omentum (mean ± s.e.m.). **f-h,** B6C3F1 mice (*n*=4/group) were treated as **a-c**. Four hours after dosing, animals were sacrificed, and tissues were analyzed ex-vivo via IVIS. Shown are weight-normalized tissue NP fluorescence (mean ± s.e.m., **f**), Pearson’s correlation coefficient for groups with significant (p<0.05) correlation between weight-normalized tissue NP fluorescence and BLI 4 hours after dosing **(g),** and representative omentum and UGT tissue IVIS BLI and NP fluorescence images for LbL NPs and UL NPs **(h)**. Correlation significance performed based on a t-test analysis with the null hypothesis of no (r=0) correlation. Group statistical comparisons in **f** were performed using a two-way ANOVA with Tukey’s multiple-comparisons test.

To define the kinetics of NP accumulation at the major sites of tumor dissemination, fluorescently labeled UL or LbL-NPs devoid of IL-12 were injected i.p. in mice bearing HM-1-luc tumors, and tumor-bearing tissues were extracted at various time points post-administration for *ex vivo* NP signal measurement. LbL-NPs showed rapid association with high tumor-burdened tissues (**Fig. 2d-e**) with peak NP fluorescence signal within ∼1 or 2 hours after injection for the UGT and omentum, respectively, at 2-5-fold higher levels than UL NPs. Evaluating Ni or Mal NPs distribution at 4 hrs post-injection showed low uptake in other organs (**Fig. 2f**) with only LbL-NPs exhibiting significant correlation between tumor bioluminescence intensity (BLI) and NP fluorescence (**Fig. 2g-h**), consistent with LbL-NP targeting of disseminated tumors.

We next examined the distribution of IL-12 in OC-bearing tissues and its spatial relationship with the NP carrier. Near the timepoint of peak NP accumulation in the UGT and omentum tissues (4 hrs), only Mal-LbL NPs exhibited a significant correlation between NP and IL-12 tissue fluorescence (**Fig. 3a**), indicating that neither the LbL film of Ni-LbL or the covalent conjugation of Mal-UL alone enabled retention of the cytokine on the particle carrier over this time course *in vivo*. At 24 hr post-dosing, animals receiving Mal-LbL NPs showed increased cytokine retention in the high tumor burdened UGT and omentum, 5- and 10-fold increased over free cytokine, respectively (**Fig. 3b**). Importantly, the fluorescence signal of IL-12 delivered via Mal-LbL NPs showed a significant correlation with tumor BLI, indicating the ability of the LbL-NPs to target disseminated OC and increase IL-12 retention in tumor tissue relative to UL or Ni NPs (**Fig. 3c**). We also assessed *ex-vivo* IVIS images for a pixel-by-pixel correlation between tumor BLI and IL-12 fluorescence. While most IL-12 treatments had a negative correlation, Mal-LbL NP delivery of the cytokine showed a positive correlation between tumor BLI and IL-12 fluorescence, suggesting improved IL-12 tumor targeting and retention with this construct (**Fig. 3d-e).**

**Figure 3.**
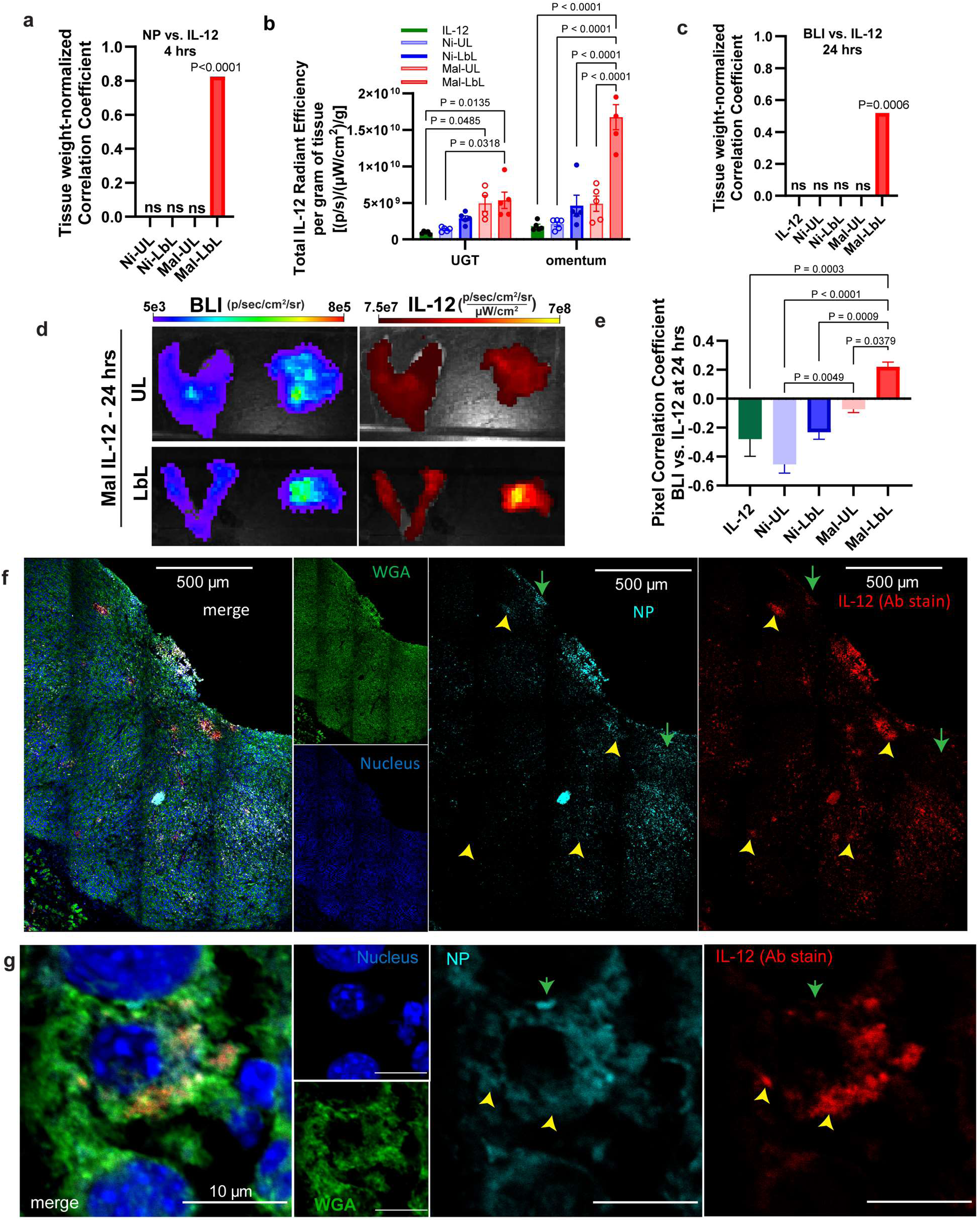
Mal-LbL NPs efficiently target and deliver IL-12 to ovarian cancer tumor nodules. **A-e,** B6C3F1 mice (n=4-5/group) inoculated with 10^6^ HM-1-luc tumor cells on day 0 were administered fluorescently-tagged NPs carrying 20 µg IL-12 on day 14. Four hours or one day after dosing, animals were sacrificed, and tissues were analyzed ex-vivo via IVIS. Shown are Pearson’s correlation coefficient for groups with significant (p<0.05) correlation between weight-normalized tissue NP fluorescence and IL-12 fluorescence 4 hours after dosing **(a)**, weight-normalized tissue IL-12 fluorescence one day after dosing in UGT and omentum (mean ± s.e.m., **b**), Pearson’s correlation coefficient for groups with significant (p<0.05) correlation between weight-normalized tissue IL-12 fluorescence and BLI one day after dosing **(c)**, representative omentum and UGT tissue IVIS BLI and IL-12 fluorescence images for Mal-UL and Mal-LbL **(d)**, and pixel-by-pixel Spearman’s correlation coefficient between IL-12 fluorescence and BLI one day after dosing from IVIS images. **E-f,** B6C3F1 mice were treated as in **a**. One day after dosing, Mal-LbL NP animals were sacrificed, and the omentum containing tumor nodules was frozen in optimal cutting temperature (OCT) compound then frozen sectioned and stained for confocal microscopy analysis. Shown are representative confocal microscopy images of tumor nodules in omental tissue at low **(f)** and high magnification **(g)**. Green arrows indicate areas with high NP signal relative to IL-12 whereas yellow arrowheads indicate areas with high IL-12 relative to NP. Correlation significance was performed based on a t-test analysis with the null hypothesis of no (r=0) correlation. Group statistical comparisons were performed using two-way ANOVA for **b** and one-way ANOVA for **e** with Tukey’s multiple-comparisons test.

Histological analysis of omental tumor nodules 24 hours after dosing revealed that the Mal-LbL NPs could efficiently penetrate tumor tissue and disseminate IL-12 (**Fig. 3f**). Consistent with the expected lipid-IL-12-conjugate release from Mal-LbL, while NP and IL-12 signals were mostly correlated across the tissue, we could observe regions with only IL-12 or only NP signal. Furthermore, at high magnification we could observe that the Mal-LbL NPs appeared to diffusely stain the membrane of cells in the tumor with patches of high IL-12 or high NP signal (**Fig. 3g**). These segregating signals suggest the release of IL-12 from the NP cores over time, on a timescale substantially slower than the time required for the NPs to effectively localize to tumor nodules.

### Tumor-targeted delivery of IL-12 is non-toxic and enhances therapeutic responses against metastatic ovarian cancer

To assess the therapeutic impact of enhanced IL-12 targeting to ovarian tumors achieved by Mal-LbL NPs, HM-1-luc tumor-bearing mice were treated with 20 µg of IL-12 administered as free cytokine or nanoparticle formulations at days 7 and 14, or a 5X higher dose of the free cytokine (to determine whether higher dosing of free cytokine could compensate for its rapid clearance, **Fig. 4a**). Three days following the first dose, a dramatic drop in tumor BLI was observed for all IL-12 treatments (**Fig. 4b**). However, except for the Mal-LbL NP-treated group, tumor signals began to rebound by day 24, ultimately leading to similar increases in median survival of ∼33 days, compared to 23 days for the untreated tumors (**Fig. 4c**). By contrast, Mal-LbL NPs delayed tumor recurrence, increasing the median survival to 44 days, with ∼30% of the animals achieving complete tumor clearance. IFN-γ ELISPOT analysis of peripheral blood lymphocytes on day 30 revealed a stronger tumor-specific T cell response induced by Mal-LbL NP treatment compared to all other groups (**Fig. 4d**). When mice that rejected their tumors were rechallenged i.p. with fresh tumor cells on day 100, all animals showed rapid tumor clearance (**Fig. S5a**) and survived whereas all naïve mice succumbed to the tumor challenge (**Fig. S5b**), indicating successful development of protective anti-tumor memory.

**Figure 4.**
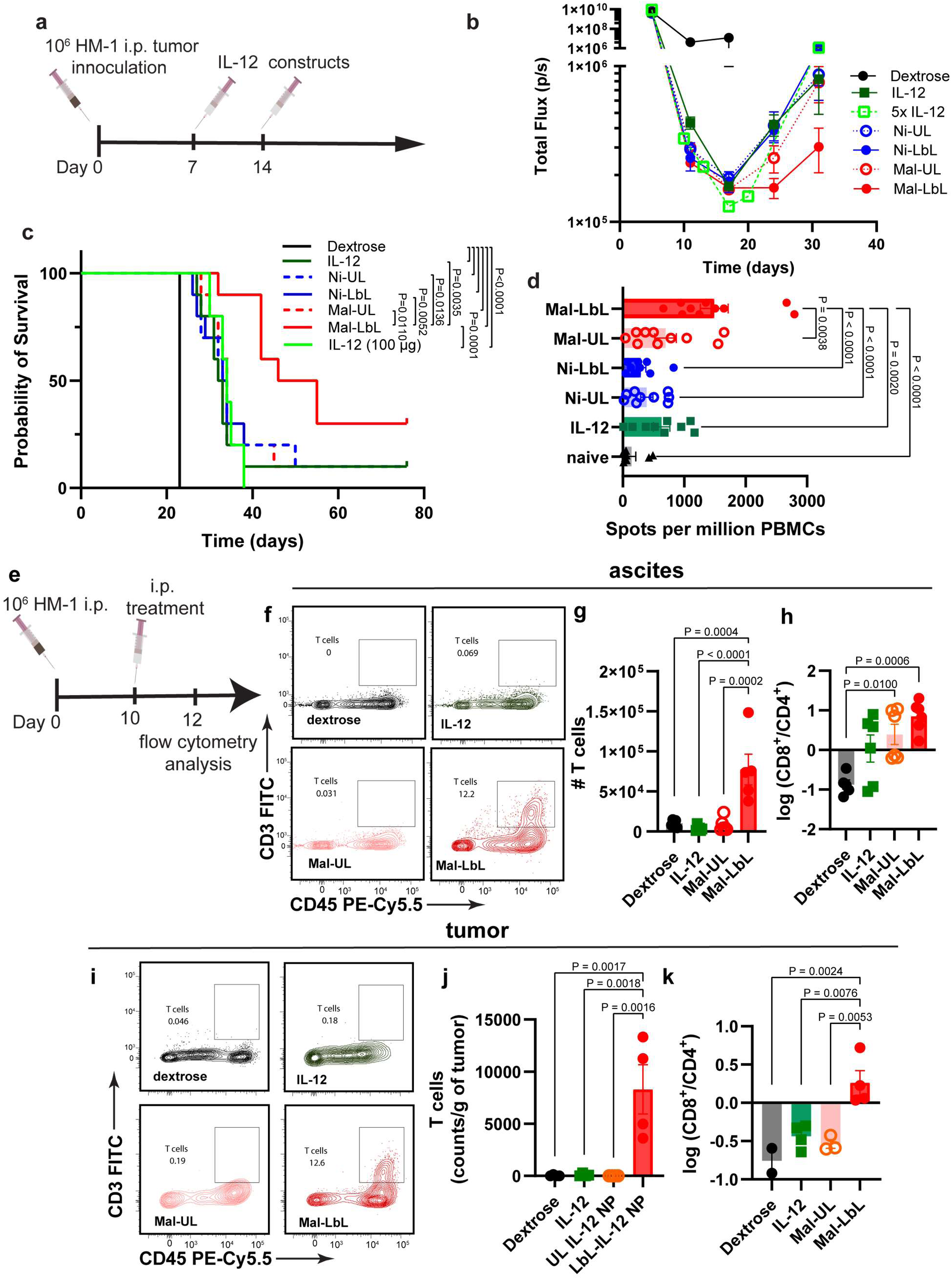
IL-12 delivered by Mal-LbL NPs exhibits potent anti-tumor activity and enhances T cell infiltration of tumors. **a-d,** B6C3F1 mice (*n* = 10/group) inoculated with 10^6^ HM-1-luc tumor cells on day 0 were treated on days 7 and 14 with 20 µg of IL-12 as a free cytokine or conjugated to NPs. Shown are the experimental timeline **(a),** in vivo IVIS whole-animal i.p. BLI readings (mean ± s.e.m., **b)**, and overall survival **(c)**. On day 30, peripheral blood mononuclear cells (PBMCs) of surviving and naïve mice (n = 5) were analyzed via IFN-γ ELISpot restimulated with HM-1-luc tumor cells. Shown are quantitation of spots detected (mean ± s.e.m., **d**). **e-k**, B6C3F1 mice inoculated with 10^6^ HM-1 tumor cells on day 0 were treated on days 10 with 20 µg of IL-12 as a free cytokine or conjugated to Mal NPs (UL and LbL). Two days after dosing ascites (*n* = 6/group) and i.p. tumor nodules (primarily omentum tissue, *n* = 4/group) were harvested and processed for flow cytometry analysis. Show are timeline for experiment (**e**), representative flow plots of T cell (CD45^+^CD3^+^) in ascites fluid (**f**), quantitation of T cells in ascites fluid (**g**), quantitation of CD8^+^ to CD4^+^ T cell ratio in ascites fluid (**h**), representative flow plots of T cell (CD45^+^CD3^+^) in tumor nodules (**i**), quantitation of T cells in tumor nodules (**j**), quantitation of CD8^+^ to CD4^+^ T cell ratio in tumor nodules (**k**). *P* values were determined by the log-rank (Mantel–Cox) test **(c)** and one-way ANOVA followed by Tukey’s multiple comparison test (**d, g, h, j, k**).

We next sought to determine the safety of this treatment. Mice bearing 10-day old HM-1 i.p. tumors were dosed with 20 µg IL-12 in either free form or conjugated to Mal-UL or Mal-LbL NPs. Two days after dosing, blood was collected, and spleen was processed via flow cytometry (**Fig. 4e**). Systemic IL-12 administration is known to increase markers of liver toxicity, induce transient cytopenia, and alter the splenic immune cell profile.^28–34^ While none of the treatments caused major liver damage as measured by liver enzyme activities, Mal-UL increased levels compared to healthy controls (**Fig. S6a**). Both free IL-12 and Mal-UL reduced white blood cell (WBC) counts compared to vehicle control mice, which was not observed with Mal-LbL treatment (**Fig. S6b)**. Untargeted IL-12 treatments also induced reduced leukocyte counts in the spleen and Mal-UL NPs elevated levels of splenic macrophage and NK cells, consistent with the expected effects of systemic IL-12 exposure (**Fig. S6d-e**).

To examine the effects of IL-12 therapy on the immune response in the local tissue, we next analyzed leukocytes in the ascites (peritoneal fluid) and tumor nodules via flow cytometry two days following a single dose of free IL-12 or IL-12 NPs at day 10 post HM-1 inoculation (**Fig. 4e**). Within the ascites, all IL-12 treatments depleted protumorigenic CD206^+^CD80^-^ (M2-like) macrophages (**Fig. S7a**), shifting the macrophage population towards a predominantly tumoricidal CD206^-^CD80^+^ M1-like phenotype (**Fig. S7b**). Polymorphonuclear and monocyte-related myeloid-derived suppressor cells (MDSCs) in the ascites, which can hinder the development of an effective immune response,^35^ were also reduced for all IL-12 treatment groups (**Fig. S7c-d**), while levels of natural killer (NK) cells increased (**Fig. S7e**). However, only Mal-LbL treatment substantially increased T cell accumulation in the ascites fluid (**Fig. 4f-g**). Characterization of the T-cell subtypes revealed a shift towards an increased CD8:CD4 ratio (**Fig. 4h**) which is associated with improved outcomes in human OC patients.^36^

In extracted tumor nodules, IL-12 treatments did not cause major effects in either PMN-MDSCs (**Fig. S7f**) or M-MDSCs levels (**Fig. S7g**). All IL-12-based treatments polarized the macrophage population from a predominantly M2-like towards a predominantly M1-like phenotype (**Fig. S7h**). However, only free IL-12 treatment led to a substantial increase in the number of M1-like tumoricidal macrophages (**Fig S7i**), suggesting a bias towards monocyte-driven immune response from systemic IL-12 treatment. On the other hand, Mal-LbL NPs increased NK cell infiltration (**Fig. S7j**). Moreover, Mal-LbL NP treatment triggered a dramatic∼50-fold increase in T cell infiltration in tumor nodules and increase in the ratio of CD8^+^ to CD4^+^ T cells, which was not observed for free IL-12 or unlayered particles (**Fig. 4i-k**). These results demonstrate the importance of targeting the cytokine to cancer cells to modulate immune infiltration of lymphocytes into tumors which is not achievable by free cytokine administration.

### Efficacy of tumor-targeted IL-12 delivery from LbL-NPs is dependent on a fluid membrane composition of the liposomal core

Membrane lipids *in vivo* can be extracted from bilayers by albumin and other serum components and undergo constant exchange with serum lipids.^37–40^ We hypothesized that the gradual release of IL-12-lipid-conjugates from Mal-LbL NPs through this process was important for optimal cytokine activity, as this could promote dissemination of the cytokine throughout the tumor bed to engage with immune cells, while simultaneously promoting prolonged retention in the tumor via insertion of the lipid tails of the conjugate in cell membranes in the local microenvironment (**Fig. 5a**). To test if IL-12-lipid conjugate release from the NP core was important for the therapeutic efficacy of Mal-LbL particles, we prepared NPs incorporating exclusively saturated lipids. Unlike Mal-LbL which contained unsaturated lipids, fully saturated lipids exhibit increased resistance to extraction by serum components *in vivo* due to their gel-phase state at physiologic temperatures (**Table S2, Fig. S8a**).^41–43^ These fully saturated LbL-NPs (SAT-LbL) carrying covalently-linked IL-12 had the same size and zeta potential as Mal-LbL particles, showed identical binding to HM-1 cells *in vitro*, and maintained IL-12 bioactivity against reporter cells (**Fig. S8b-f)**. However, SAT-LbL NPs were substantially more resistant to the extraction of fluorophore tagged lipids in serum compared to Mal-LbL (**Fig. S8g)** and showed reduced IL-12 release in serum compared to Mal-LbL NPs (**Fig. S8h)**.

**Figure 5.**
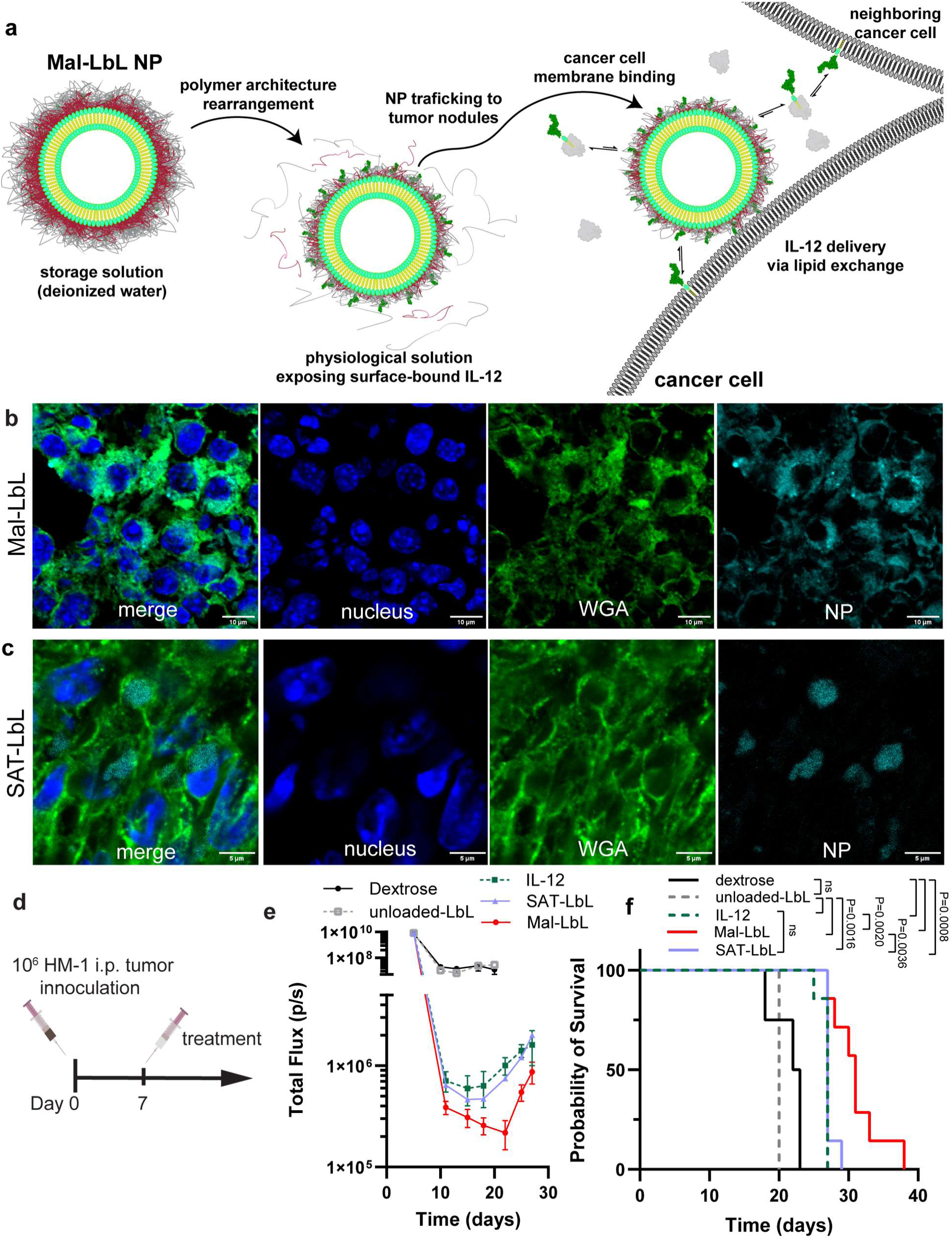
Efficacy of Mal-LbL NPs is dependent on lipid exchange properties which alter lipid distribution inside tumor nodules. **a,** Schematic of proposed mechanism of tumor targeted IL-12 lipid-conjugate dissemination from Mal-LbL NPs. **b-c**, B6C3F1 mice were inoculated with 10^6^ HM-1-luc tumor cells on day 0 were administered fluorescently-tagged Mal-LbL of SAT-LbL NPs carrying 20 µg IL-12 on day 14. One day after dosing, animals were sacrificed, and the omentum containing tumor nodules was frozen in optimal cutting temperature (OCT) compound then frozen sectioned and stained for confocal microscopy analysis. Shown are representative high-magnification confocal images of omental tumor nodules from Mal-LbL (**b**) and SAT-LbL (**c**). **d-f**, B6C3F1 mice (*n* = 7/group) inoculated with 10^6^ HM-1-luc tumor cells on day 0 were treated on days 7 with NP vehicle control (unloaded-LbL), 20 µg of IL-12 as a free cytokine or conjugated to Mal-LbL or SAT-LbL. Shown are the experimental timeline **(d),** in vivo IVIS whole-animal i.p. BLI readings (mean ± s.e.m., **e)**, and overall survival **(f)**. Statistical comparisons between survival curves were performed using a log-rank (Mantel–Cox) test.

To determine whether membrane composition impacts the biodistribution of lipids initially incorporated into the liposomal core, we treated HM-1 tumor-bearing mice with either Mal-LbL or SAT-LbL NPs carrying fluorescent phosphatidylethanolamine (PE) lipids. One day after dosing, we extracted the omentum tumor nodules for histological assessment via confocal microscopy. Both SAT-LbL and Mal-LbL NPs efficiently penetrated tumors (**Fig. S9a-b**). Strikingly, the membrane composition of the NPs directly impacted the pattern of fluorescent-lipid distribution in tumor nodules: fluorophore-lipids delivered by Mal-LbL NPs localized in regions showing high staining for cell membranes and extracellular matrix (wheat germ agglutinin, WGA), while fluorescent lipids delivered as part of SAT-LbL NPs localized in pockets devoid of WGA staining **(Fig. 5b-c**), suggesting that SAT-LbL particles (and their associated fluorescent-lipid cargo) might be ultimately internalized by cells in the tissue.

We next carried out a therapeutic study administering a single dose of 20 µg IL-12 as free cytokine, Mal-LbL NPs, SAT-LbL NPs, or a control LbL-NP lacking IL-12 (unloaded-LbL, **Fig. 5d**). Unloaded-LbL NPs had no impact on tumor BLI and did not induce any therapeutic benefit compared to vehicle control mice (**Fig. 5e-f**). As seen in the two-dose setting, a single administration of Mal-LbL NPs dramatically reduced tumor BLI by 3 days post dosing, which did not begin to relapse until after day 22 and extended median survival to 31 days. Interestingly, free IL-12 and SAT-LbL NPs elicited very similar tumor BLI reduction/relapse and overall survival, with relapse evidence after day 18 and median survival of only 27 days (**Fig. 5e-f**). These data suggest that the lipid-exchange properties of Mal-LbL NPs enabling slow release of lipid-conjugate IL-12 are important for the improved therapeutic benefit seen with this formulation.

### Mal-LbL NPs sensitize metastatic ovarian cancer to immune checkpoint inhibitors therapy

Checkpoint inhibitors (CPI), such as antibodies blocking the negative regulatory receptors PD-1 and CTLA-4 expressed by T cells, are currently the most broadly effective class of immunotherapy agents clinically, but CPIs have failed to show substantial benefit in OC.^44,45^ Responsiveness to checkpoint blockade correlates with the presence of pre-existing CD8 T cell infiltrates in patient tumors,^46^ and thus we hypothesized that enhanced T cell infiltration driven by LbL-NPs could increase the responsiveness of OC to checkpoint inhibition. Moreover, IL-12 induces strong IFN-γ expression in T cells and NK cells, which will in turn upregulate expression of PD-L1 on tumor cells, a phenomenon termed adaptive resistance.^47^ We thus theorized that Mal-LbL treatment may sensitize OC to CPI therapy. To test this idea, we evaluated treatment of HM-1-luc tumors with systemic anti-PD-1 + anti-CTLA-4 CPI therapy alone or combined with two weekly doses of IL-12 in free or NP form (**Fig. 6a**). Analysis of tumor burden via BLI clearly showed that CPIs alone could only mildly and transiently control tumor growth (**Fig. 6b**). On the other hand, there was a marked synergism when any form of IL-12 was included in the treatment, reducing tumor BLI to baseline for one month after tumor inoculation. CPI therapy alone had only marginal survival benefit over untreated controls, similar to what has been observed with OC in the clinic (**Fig. 6c).**^45^ Combining CPI with free IL-12 showed some efficacy, reducing tumor BLI to baseline for ∼25 days and curing 20% of treated animals (**Fig. 6b-c**). Ni-LbL also synergized with CPI treatment, showing a significant extension in survival compared to free IL-12 and CPI but ultimately resulted in only a 30% long-term survivor rate. Strikingly, however, Mal-LbL showed a remarkable sensitization effect, achieving 100% cures with this treatment schedule (**Fig. 6c**). When challenged with fresh tumor cells at day 150, all Mal-LbL NP-treated mice rejected the rechallenge (**Fig. 6d-e**), demonstrating induction of a strong memory response. Thus, engineering the rapid targeting and slow local release of IL-12 in ovarian tumor nodules using LbL-NPs dramatically sensitizes tumors to clinically approved CPI combination therapy.

**Figure 6.**
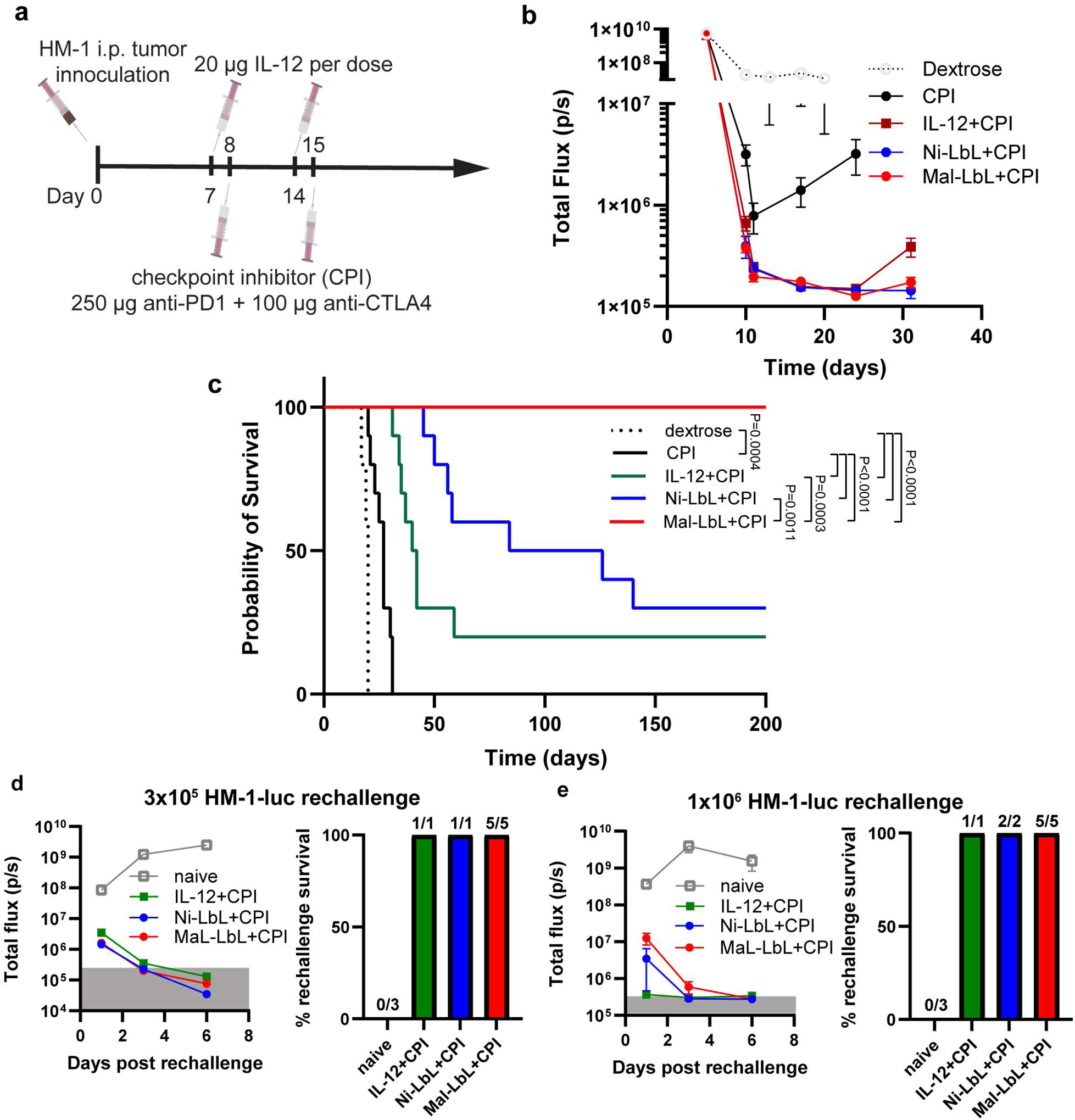
Combination of immune checkpoint inhibitors with two dose treatment of Mal-LbL eradicates metastatic ovarian cancer. **a-e,** B6C3F1 mice (*n* = 10/group) inoculated with 10^6^ HM-1-luc tumor cells on day 0 were treated on days 7 and 14 with 20 µg of IL-12 as a free cytokine or conjugated to Mal-LbL or Ni-LbL. Mice were also treated with 250 µg of anti-PD1 and 100 µg of anti-CTLA4 i.p. on days 8 and 15. Shown are the experimental timeline **(a),** *in vivo* IVIS whole-animal i.p. BLI readings (mean ± s.e.m., **b)**, and overall survival **(c)**. On day 150, surviving mice were rechallenged with either 3×10^5^ (**d**) or 10^6^ (**e**) HM-1-luc tumor cells i.p. Shown are in vivo IVIS whole-animal i.p. BLI readings post rechallenge (mean ± s.e.m) and the percentage of mice per group that survived rechallenge. Statistical comparisons between survival curves were performed using a log-rank (Mantel–Cox) test.

## Discussion

Cytokine therapies have been challenging to develop and achieve regulatory approval.^9,48^ IL-12 has long been viewed as an attractive candidate cytokine to drive ant-tumor immunity, but has failed to progress clinically due to its low therapeutic index when administered as a free protein drug.^49^ Here we report the use of LbL-NPs capable of targeting and attaching to cancer cell membranes to deliver IL-12 in peritoneally-disseminated metastatic OC. We demonstrate that optimized LbL-IL-12-NPs are non-toxic, elicit strong systemic anti-tumor immunity, drive remodeling of the TME, and strongly sensitize ovarian tumors to CPI therapy. Importantly, we demonstrate that the efficacy of LbL-IL-12-NPs is dependent on both covalent conjugation of the cytokine payload to lipids and lipid extraction, a known but relatively unexplored property of lipid-based nanoparticles. It is also clear that targeting the LbL-NPs to the tumor cell membrane surface is a critical aspect of this approach. In addition to delayed lipid-conjugate release from the NP, such a mechanism may promote trafficking of therapeutic payloads to tumor-draining lymph nodes.^50^

Although other approaches exist to augment cytokine delivery in cancer, most rely on local (intratumoral, i.t.) administration to increase tumor drug concentration while minimizing systemic exposure.^51^ However, such approaches would be impractical to implement in metastatic OC due to difficulties with administration into small, disseminated legions in the i.p. space. Moreover, while NPs have previously been explored for cytokine delivery, they have primarily relied on passive tumor targeting through the enhanced permeability and retention (EPR) effect by extending NP circulatory half-life via polyethylene glycol (PEG) decoration.^8^ However, in addition to concerns with anti-PEG immune responses^52^, increased half-life also increases systemic exposure and subsequent toxicity and the lack of specific tumor interactions would facilitate clearance from the peritoneal cavity after i.p. administration.^17^

In summary, this study demonstrates that engineering PLE-coated LbL-NPs carrying IL-12 can facilitate both tumor-targeted delivery and sustained cytokine release within disseminated tumor nodules. Combining tumor targeting with localized cytokine dissemination in the tumor microenvironment significantly improves the efficacy of this immunotherapy against metastatic ovarian cancer. Importantly, optimized LbL-NPs exhibited strong synergy with checkpoint blockade, the current gold standard for cancer immunotherapy in the clinic.

## Methods

### Materials

1,2-distearoyl-sn-glycero-3-phosphocholine (DSPC), 1,2-dioleoyl-sn-glycero-3-[(N-(5-amino-1-carboxypentyl)iminodiacetic acid)succinyl] (nickel salt) (18:1 (Ni)NTA-DGS), 1,2-dioleoyl-sn-glycero-3-phosphoethanolamine-N-[4-(p-maleimidophenyl)butyramide] (sodium salt) (18:1 MPB-PE), 1-palmitoyl-2-oleoyl-sn-glycero-3-phospho-(1’-rac-glycerol) (sodium salt) (POPG), 1,2-dioleoyl-sn-glycero-3-phosphoethanolamine-N-dibenzocyclooctyl (DOPE-DBCO), 1,2-distearoyl-sn-glycero-3-phospho-(1’-rac-glycerol) (sodium salt) (DSPG), 1,2-dipalmitoyl-sn-glycero-3-phosphoethanolamine-N-[4-(p-maleimidophenyl)butyramide] (sodium salt) (16:0 MPB-PE), 1,2-dipalmitoyl-sn-glycero-3-phosphoethanolamine-N-dibenzocyclooctyl (16:0 DBCO PE), and cholesterol were purchased from Avanti Polar Lipids. Poly-L-arginine (PLR) with a molecular weight (MW) of 9.6 kDa and poly-L-glutamic acid (PLE) with a MW of 15 kDa were purchased from Alamanda Polymers. BDP TMR azide (Lumiprobe) and BDP 630/650 azide (Lumiprobe) were conjugated to DOPE-DBCO or 18:0 DBCO-PE in chloroform to generate fluorescently labeled lipids. Successful conjugation was validated via thin-layer chromatography which indicated <1% free dye. For immunophenotyping (all mouse targets), antibodies for CD3e (clone 145-2C11, FITC), CD49b (clone DX5, PE), CD45 (close 30-F11, PE-Cy5.5), F4/80 (clone T45-2342, PE), CD80 (clone 16-10A1, FITC), CD11b (clone M1/70, FITC), and Ly6C (clone AL-21, PE) were purchased from BD Biosciences. Antibodies for CD4 (clone GK1.5, APC-Cy7), CD8a (clone 53-6.7, PE), CD206 (clone C068C2, PE-Cy5), and Ly6G (clone 1A8, APC) were purchased from BioLegend. Deionized water was of ultrapure grade obtained through a Milli-Q water system (EMD Millipore).

### Recombinant single-chain IL-12 production

Single-chain IL-12 sequence^53^ was synthesized as a genomic block (Integrated DNA Technologies) and cloned into gWIZ expression vector (Genlantis). Plasmids were transiently transfected into Expi293 cells (ThermoFisher Scientific). After 5 days, cell culture supernatants were collected and protein was purified in an ÄKTA pure chromatography system using HiTrap HP Niquel sepharose affinity column, followed by size exclusion using Superdex 200 Increase 10/300 GL column (GE Healthcare Life Sciences). Endotoxin levels in purified protein was measured using Endosafe Nexgen-PTS system (Charles River) and assured to be <5 EU/mg protein.

### Liposome synthesis

A lipid solution was prepared by mixing (all mole %) 65% DSPC (25 mg/mL), 24% cholesterol (25 mg/mL), 6% POPG (25 mg/mL) and 5% of either 18:1 (Ni)NTA-DGS (5 mg/mL) or 18:1 MPB-PE (5 mg/L) then formed into a thin film using a rotary evaporator (Buchi). Lipid films were allowed to further dry overnight in a desiccator, then were hydrated at 0.5-1 mg/mL using deionized water and sonicated for 3-5 minutes at 65 °C then extruded (Avestin Liposofast LF-50) once at 65 °C through a 100 nm membrane (Cytiva Nuclepore) then 3X through 50 nm membranes (Cytiva Nuclepore) . Extruded liposomes were added to an ice bath. SAT NPs were generated with the same procedure as Mal NPs, but its composition (all mole %) was 65% DSPC (25 mg/mL), 24% Cholesterol (25 mg/mL), 6% DSPG (25 mg/mL) and 5% of 16:0 MPB-PE (5 mg/mL). Lipid were stored at -20 °C in amber vials in chloroform except DSPG which was stored in a 1:1 (v/v) mixture of chloroform and methanol instead of pure chloroform.

For coupling of scIL-12 via Ni-histag interactions, scIL-12 was added to 0.5 mg/mL Ni(NTA)-DGS liposomes at a molar ratio of 28:1 of Ni(NTA)-DGS lipids to IL-12. After incubation with IL-12 at 4 °C for 18 hr, Ni-UL liposomes were purified via tangential flow filtration on a 100 kDa mPES membranes (Repligen) against 6 diafiltration volumes of deionized water.

For covalent linkage of scIL-12 to Mal-liposomes, the solution pH of MPB-PE liposomes was adjusted to pH 5 with hydrochloric acid prior to lipid film hydration, and following membrane extrusion, liposomes at 0.33 mg/mL were adjusted to pH 7.0 with 10 mM HEPES prior followed by addition of scIL-12 containing a terminal cysteine residue at a molar ratio of 25:1 of MPB-PE lipid to protein for at least 12 hours at 4 °C in a rotating mixed. Any remaining maleimides were quenched with a 100-fold molar excess of L-cysteine (Sigma) for 1.5 hrs on ice.

For fluorescence labeling of liposomes, 0.2 mol% of DSPC content was replaced by either DOPE-TMR or DOPE-630/650. IL-12 concentrations were measured via enzyme-linked immunoassay (ELISA) (Peprotech) and lipid content was quantified via the Stewart Assay.^54^

### Layer-by-Layer (LbL) film deposition onto NPs

Assembly of polyelectrolyte layers was performed by adding unlayered particles to a diH2O solution with 0.3-0.4 weight equivalents (wt.eq.) of PLR relative to lipid in a glass vial under sonication and incubating on ice for at least 30 min. Excess PLR polymer was purified by TFF through a 100 kDa mPES membrane (Repligen) pre-treated with a 10 mg/mL solution of free PLR. For the terminal PLE layer, purified particles coated with PLR were added to a diH2O solution with PLE in a glass vial under sonication at 1 wt.eq. of polymer to lipid. LbL particles were then purified by TFF on a separate 100 kDa mPES membrane (Repligen) to remove any excess PLE.

### Characterization of particle preparations

Dynamic light scattering (DLS) and zeta potential measurements were made on a Zetasizer Nano ZSP (Malvern). Nanoparticle micrographs were acquired using Transmission Electron Microscopy (TEM) on a JEOL 2100F microscope (200 kV) with a magnification range of 10,000-60,000X. All images were recorded on a Gatan 2kx2k UltraScan CCD camera. Negative-stain sample preparation was performed by adding 10 µL of NPs on a 200 meshes copper grid coated with a continuous carbon film and allowing for sample adsorption for 60 seconds. Excess solution was then removed by touching the grid with a kimwipe. The grid was then quickly washed by adding 10 µL of negative staining solution, phosphotungstic acid (PTA), 1% aqueous solution then removing excess by touching the grid with a kimwipe. Then, the grid was mounted on a JEOL single tilt holder equipped in the TEM column for image capture.

### Fluorescent labeling of polymers

PLR was solubilized at 100 mg/mL in diH2O then mixed with 2 molar equivalents of BDP-TR-NHS-ester (Lumiprobe) in DMSO to generate a 15 mg/mL PLR solution. The reaction was catalyzed with ten molar equivalent of triethylamine (TEA) and allowed to react for 4 hrs at room temperature then overnight at 4 °C. PLR-TR was purified via Reverse-Phase High Pressure Liquid Chromatography (RP-HPLC) on a Jupiter C4 column (5 µm particles, 300 Å – Phenomenex) using a water:acetonitrile gradient which started at 20% actetonitrile for 5 minutes, then increased to 35% in a linear gradient until 10 minutes. Isocratic elution at 35% was performed for 30 minutes then the elution buffer was increased to 95% to clean out the column for 10 minutes then dropped back to 20% acetonitrile to re-equilibrate the column for 5 minutes. Collected purified PLR-TR fractions were then diluted 10-fold with diH2O then lyophilized. PLE at 10 mg/mL was labeled by reacting with 5 molar equivalents of sulfo-cyanine3 NHS ester (Lumibrobe) in PBS adjusted to pH ∼8.5 with 0.1 M sodium bicarbonate. Excess dye was removed via extensive 0.9wt% NaCldialysis followed by extensive diH2O dialysis using a 3 kDa regenerated cellulose membrane (Repligen) and the purified PLE-cy3 was lyophilized until use.

### Analysis of LbL film stability

For PLR stability, PLR/PLE films were assembled onto Mal-UL NPs as described in *Layer-by-Layer (LbL) film deposition onto NPs,* but the PLR solution was doped with 30% PLR-TR. For PLE stability, the PLE solution was doped with 50% PLE-cy3. Particles were incubated in various buffer solutions at 0.1 mg/mL in a shaker at 37 °C. At defined intervals, aliquots were extracted from the incubation solution and free polymers were separated from the NP. For PLE separation, samples were spun on a 300 kDa centrifugal filter (VivaSpin500, Sartorius) at 10x*g* for 15 min and the permeate fluorescence was compared to the fluorescence of the initial sample. For PLR separation, the NPs in the sample were pelleted by centrifuging at 25000x*g* for 30 min and the supernatant PLR fluorescence was compared to the initial sample PLR fluorescence. Particles were validated to lack free polymers by centrifuging in diH2O. Fluorescence was measured on 96 well plates using a plate reader (Tecan Infinite 200).

### IL-12 accessibility via monoclonal antibody binding

IL-12 accessibility to monoclonal antibody binding was determined by using the antibodies in an IL-12 sandwich ELISA kit (Peprotech). After coating a 96 well plate with anti-IL-12 antibodies and blocking the plate with bovine serum albumin according to the manufacturer protocol, Mal-UL or Mal-LbL NPs were captured onto the plate by incubating the samples for two hours in either diH2O or HEPES buffered saline solution supplemented with 10% FBS. After sample incubation with capture antibodies, plates were washed and total captured IL-12 was determined following the manufacturer’s protocol.

### Cell Culture

OV2944-HM-1 cells were acquired through Riken BRC and were cultured in α-MEM. HEK-Blue IL-12 (InvivoGen) cells were cultured and used for IL-12 bioactivity assessment according to the manufacturer’s instructions. Cell media was also supplemented with 10% FBS and penicillin/streptomycin with cells incubated in a 5% carbon dioxide humidified atmosphere at 37°C. All cell lines were murine pathogen tested and confirmed mycoplasma negative by Lonza MycoAlert™ Mycoplasma Detection Kit.

### In vitro cellular association

HM-1 cells were plated on a tissue-culture 96-well plate at a density of 50K cells per well. The next day, wells were dosed with NPs and left for the target incubation time (4 hrs or 24 hrs). For analysis via flow cytometry, NPs were dosed at 0.02 mg/mL and allowed to incubate with cells for 4 hours at 37°C. Cells were washed with PBS then detached from the plates using 0.25% trypsin and stained with DAPI (15 min incubation) for viability assessment and fixed with 2% paraformaldehyde (30 min incubation) until analysis by flow cytometry using an LSR Fortessa (BD Biosciences). For assessment of NP-associated fluorescence in a fluorescence plate reader, a dose of 0.05 mg/mL was used and before cell washing with PBS, the supernatant was removed from the well and diluted 10X with DMSO. Cells were then washed three times with PBS and disrupted with DMSO. Fluorescence of NPs associated with cells was then normalized to supernatant fluorescence. The relative fluorescence of each formulation was then compared to an unlayered liposome control containing the same fluorophore. For confocal imaging, 8-well chambered coverglasses (Nunc Lab-Tek II, Thermo Scientific) were coated with rat tail collagen type I (Sigma-Aldrich) per the manufacturer’s instructions. HM-1 cells were plated into the wells at a density of 10K/well and left to adhere overnight prior to NP treatment. After the desired incubation time with NPs, cells were washed 3x with PBS. After washing, cells were fixed in 4% paraformaldehyde for 10 minutes then washed (3x with PBS) and stained with wheat germ agglutinin (WGA) conjugated to Alexa Fluor488 (Invitrogen) and Hoechst 33342 (Thermo Scientific) following manufacturer’s instructions. Images were analyzed using ImageJ. Slides were imaged on an Olympus FV1200 Laser Scanning Confocal Microscope.

### Fluorescent labeling of IL-12

IL-12 was labeled with indocyanine green (ICG) tetrafluorophenyl (TFP) ester (AAT Bioquest) by solubilizing the dye in dimethyl sulfoxide at 1 mg/mL and adding it to IL-12 at 3 mg/mL in phosphate buffered saline (PBS) supplemented with 0.1 M sodium bicarbonate at a 1.2:1 molar ratio of dye to protein. Excess dye was removed via 7 kDa desalting columns (Zeba Spin, ThermoFisher) and validated via thin-layer chromatography.

### IL-12 release assay

IL-12 release from liposomes was quantified using the same procedure as quantification of PLE stability.

### Serum-induced lipid exchange

Lipid release from NPs was assessed by generating liposomes with a high density (1 mol%) of DOPE-630-650 (for Mal-LbL) or DPPE-630/650 (for SAT-LbL) to induce fluorescence quenching. Particles were then mixed with 100% FBS solution supplemented with penicillin/streptomycin and incubated in 96 well plates in a shaker at 37 °C. At certain intervals, dye fluorescence was measured (ex: 610 nm/ em: 650 nm) on a plate reader (Tecan Infinite200) and compared to total dye fluorescence which was obtained by dissolving the NPs in DMSO.

### Animals

All animal work was conducted under the approval of the Massachusetts Institute of Technology Division of Comparative Medicine IACUC in accordance with federal, state, and local guidelines. B6C3F1 mice were purchased from Jackson Laboratories. Female mice were used between 8-12 weeks of age unless otherwise noted.

### Kinetics of NP association with high tumor burden tissues in metastatic ovarian cancer model

B6C3F1 mice were inoculated with firefly luciferase-expressing OV2944-HM1 (HM-1-luc) cells through intraperitoneal (i.p.) injection of 10^6^ cells in PBS. For kinetics of LbL-NP association, two weeks after tumor inoculation, mice were injected with NPs containing fluorescently labeled lipid (DOPE-630/650, 0.2 mol%) and euthanized 1, 2, 4, 12 and 24 hrs after dosing. UGT and omentum tissues were extracted and placed in PBS under ice. Tissues were then transferred to an In Vivo Imaging System (IVIS, Perkin Elmer) to quantify ex-vivo tissue NP fluorescence. Data were analyzed using Living Image software. Background fluorescence measurements were made for each organ based on signal from mice treated with dextrose and measurements were normalized to tissue weight.

### Pharmacokinetic and biodistribution in metastatic ovarian cancer model

B6C3F1 mice were inoculated with firefly luciferase-expressing OV2944-HM1 (HM-1-luc) cells through intraperitoneal (i.p.) injection of 10^6^ cells in PBS. Two weeks after tumor inoculation, mice were injected with NPs containing fluorescently labeled lipid (DOPE-630/650, 0.2 mol%) and IL-12-ICG. The same IL-12-ICG was used for all groups to avoid labeling efficiency differences and groups were dosed intraperitoneally with 20 µg of IL-12. *In vivo* tumor radiant efficiency was measured on an IVIS by imaging the mice i.p. region. After the final timepoint (4 hrs or 24 hrs), mice were euthanized, organs were removed and placed in PBS under ice. Organs were then transferred to a 1 mg/mL D-luciferin solution in PBS and incubated for 5 minutes then placed in an IVIS to determine each organ’s BLI, NP fluorescence and IL-12 fluorescence. Data were analyzed using Living Image software. Background fluorescence measurements were made for each organ based on signal from mice treated with dextrose. For correlation analysis, the weight-normalized bioluminescence flux (p/s/g) and radiant efficiency ([p/s] / [µW/cm²]/g) for each organ were analyzed on Graphpad Prism 9 for their correlation via the Pearson’s correlation coefficient.

### IVIS image pixel correlation analysis

IVIS images were extracted from the Living Image software using the same range of BLI and IL-12 fluorescence values. Pixel intensity values for all tissues of each mouse were extracted using ImageJ and then analyzed for Spearman’s correlation between BLI and IL-12 using GraphPad Prism 9.

### Cryogenic freezing of omentum tumors

Omentum tissue from the biodistribution study was added to optimal cutting temperature (OCT) compound and rapidly frozen in cryomolds using isopentane with dry ice. Samples were sectioned in 10 µm slices on a microtome-cryostat onto Tissue Path Superfrost Plus Gold Slides (Fisherbrand) and stored in -80 °C. For staining, slides were rapidly fixed with ice-cold 4% methanol-free formaldehyde for 10 minutes then washed with PBS and blocked with 10% goat serum for 1 hr. The samples were then incubated with PE anti-IL-12/IL-23 p40 antibody (Biolegend) overnight at 4 °C in 1% bovine serum albumin (BSA) PBS buffer. After overnight incubation, Hoechst 33342 (ThermoFisher) and WGA-alexafluor488 (ThermoFisher) were added and allowed to incubate for 30 minutes at room temperature. Samples were then washed with PBS then were mounted with a coverslip using ProLong Gold (ThermoFisher) and stored at 4 °C after drying. Slides were imaged on an Olympus FV1200 Laser Scanning Confocal Microscope.

### Immunophenotyping via flow cytometry and blood panel analysis

B6C3F1 mice were inoculated intraperitoneally with 10^6^ cells of HM-1 in PBS. Ten days after tumor inoculation, mice were treated with either dextrose (vehicle control) or 20 µg of IL-12 in free, Mal-UL, or Mal-LbL formats. Two days after dosing, mice were bled retro-orbitally and then euthanized to extract ascites cells via peritoneal lavage with PBS. Peritoneal tumor nodules and spleen were also collected. Part of the blood samples were submitted to The Division of Comparative Medicine at MIT to perform a complete blood count and analysis of liver function and the remainder was processed with ACK lysing buffer (Gibco) to isolate PBMCs. Spleens were processed on a 70 µm cell strainer with a syringe plunger then exposed to ACK lysing buffer to lyse red blood cells (RBCs). Tumor nodules from each mouse were diced with scissors then incubated for one hour at 37 °C in 2 mL of 1 mg/mL collagenase type IV (Sigma) in RPMI media. After collagenase incubation, tumors were processed on a 70 µm cell strainer then collected with an insulin syringe into falcon tubes to pellet tumor cells and wash out collagenase solution. For cell staining with antibodies, samples were placed in 96 well plates, then centrifuged and resuspended in Fc block solution (BD Biosciences) for 5 minutes. Freshly prepared antibody panels were then mixed with the samples and allowed to react for 20 minutes. Finally, DAPI (BD Biosciences) was added to each well (at 2 µg/mL) and allowed to react for 5 minutes. Stained cells were washed twice with flow cytometry buffer (PBS with 0.5% BSA and 2 mM EDTA), then resuspended in 2% PFA in PBS for 30 minutes, washed and stored at 4 °C for analysis the next day on a on a flow cytometry instrument (LSR Fortessa, BD Biosciences). Flow cytometry buffer was used to prepare Fc block and antibody solutions. The gating strategy for flow cytometry analysis with each antibody used is shown in **Fig. S10**.

#### Efficacy studies with metastatic ovarian cancer model

B6C3F1 mice were inoculated intraperitoneally with 10^6^ cells of HM-1-luc in PBS. One week after inoculation, treatment was initiated as indicated on each figure. All treatments included the same IL-12 dose. For combination with immune checkpoint inhibitors, mice received 250 µg of anti-PD1 antibody (clone 29F.1A12, BioXCell) and 100 µg of anti-CLTA4 antibody (clone 9D9, BioXCell) i.p. one day after treatment with IL-12 constructs. Animal weights were tracked daily after treatments for signs of toxicity. Bioluminescence was measured on a IVIS 10 minutes after i.p. injection of 3 µg of D-luciferin sodium salt (GoldBio) for 30 days after tumor inoculation or as needed to track tumor burden.

#### IFN-γ ELISPOT

Blood was collected from mice via sub-mandibular bleeding and lysed in ACK Lysis Buffer then placed in RPMI supplemented with 10% FBS, 1% penicillin–streptomycin, 1× non-essential amino acids (Invitrogen), 1× sodium pyruvate (Invitrogen) and 1× 2-mercaptoethanol (Invitrogen). On the same day, HM-1-luc cells (treated with 500 U ml−1 IFN-γ overnight) were subjected to 120 Gy radiation and trypsinized into a single-cell suspension in the same supplemented RPMI. Then, 25,000 irradiated HM-1-luc cells were mixed with 3 × 10^5^ PBMCs per sample and seeded in a 96-well ELISPOT plate (BD Biosciences) that was pre-coated with IFN-γ capture antibody (BD Biosciences). Plates were cultured for 24 h in a 37 °C incubator, then developed according to the manufacturer’s protocol. Plates were scanned using a CTL-ImmunoSpot Plate Reader, and data were analysed using CTL ImmunoSpot Software.

#### Statistical Analysis

GraphPad PRISM 9 was used to perform statistical analyses. Comparisons between two groups was performed via unpaired t-tests. For multiple groups or multiple variable analysis, one-way, or two-way ANOVAs were used with Tukey’s posthoc correction for time-based analysis or Sidak posthoc for other ANOVA analysis.

## Acknowledgments

We thank the Koch Institute Swanson Biotechnology Center for technical support.

## Disclosure Statement

ISP, PTH, and DJI are inventors on a patent filed by the Massachusetts Institute of Technology relating to LbL NP therapeutics.

## Funding Sources

This work was supported in part by the National Institutes of Health (awards R01CA235375 to PTH and DJI, and F99CA274651 to ISP), the Marble Center for Nanomedicine, and the Ragon Institute of MGH, MIT, and Harvard. DJI is an investigator of the Howard Hughes Medical Institute. This work was also supported by the Koch Institute Support (core) Grant P30-CA14051 from the National Cancer Institute.

## Data availability

Raw data are available from the corresponding author upon request.

