## Supplementary Information for "“Target-and-release” nanoparticles for effective immunotherapy of metastatic ovarian cancer"

**Table S1: Lipid composition of nanoparticles and summary characteristics.**

| **Lipid or trait** | **Nickel-headgroup** | **Maleimide-headgroup** |
| --- | --- | --- |
| DSPC; 1,2-distearoyl-sn-glycero-3-phosphocholine | 65 mol% | 65 mol% |
| Cholesterol | 23.9 mol% | 23.9 mol% |
| POPG; 1-palmitoyl-2-oleoyl-sn-glycero-3-phospho-(1'-rac-glycerol) (sodium salt) | 6.1 mol% | 6.1 mol% |
| 18:1 DGS-NTA(Ni); 1,2-dioleoyl-sn-glycero-3-[(N-(5-amino-1-carboxypentyl)iminodiacetic acid)succinyl] (nickel salt) | 5 mol% | 0 mol% |
| 18:1 MPB-PE; 1,2-dioleoyl-sn-glycero-3-phosphoethanolamine-N-[4-(p-maleimidophenyl)butyramide] (sodium salt) | 0 mol% | 5 mol% |
| Diameter (Z-avg) unlayered (UL) | 86.3 nm | 87.7 nm |
| Diameter (#-avg) unlayered (UL) | 60.2 nm | 57.8 nm |
| PDI unlayered (UL) | 0.08 | 0.14 |
| Zeta potential unlayered (UL) | -58 mV | -63 mV |
| Diameter (Z-avg) PLR-PLE (LbL) | 121.3 nm | 122.5 nm |
| Diameter (#-avg) PLR-PLE (LbL) | 84.7 nm | 81.2 nm |
| Zeta potential PLR-PLE (LbL) | -63 mV | -63 mV |
| PDI PLR-PLE (LbL) | 0.11 | 0.13 |

**Table S2: Lipid composition of SAT NPs.**

| **Lipid** | **SAT NPs**  **(mol%)** |
| --- | --- |
| DSPC; 1,2-distearoyl-sn-glycero-3-phosphocholine | 65% |
| Cholesterol | 23.9% |
| DSPG; 1,2-distearoyl-sn-glycero-3-phospho-(1'-rac-glycerol) (sodium salt) | 6.1% |
| 16:0 MPB-PE; 1,2-dipalmitoyl-sn-glycero-3-phosphoethanolamine-N-[4-(p-maleimidophenyl)butyramide] (sodium salt) | 5% |

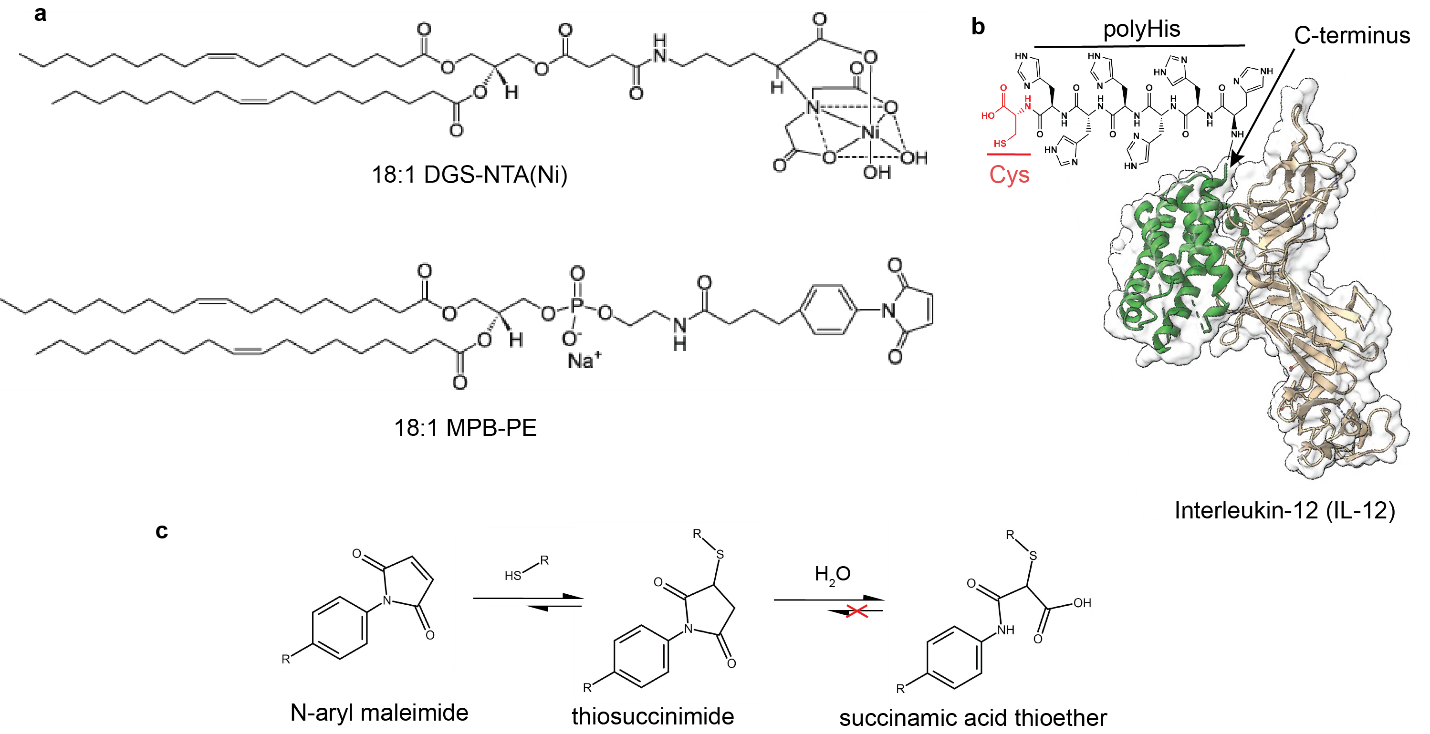

**Figure S1.** **Headgroup-modified lipids for IL-12 conjugation. a,** Chemical structure of headgroup-modified lipids with either chelated nickel or N-aryl maleimide. N-aryl maleimide was employed to prevent potential thiosuccinimide retro-Michael addition and subsequent thiol-exchange, as this headgroup favors thiosuccinimide hydrolysis into stable succinamic acid thioethers. **b**, IL-12 crystal structure derived from PDB 1F45 showing C-terminus used to engineered terminal poly-histidine (polyHis) tag and terminal cysteine. **c,** Reaction pathway for irreversible maleimide-based conjugation with thiols.

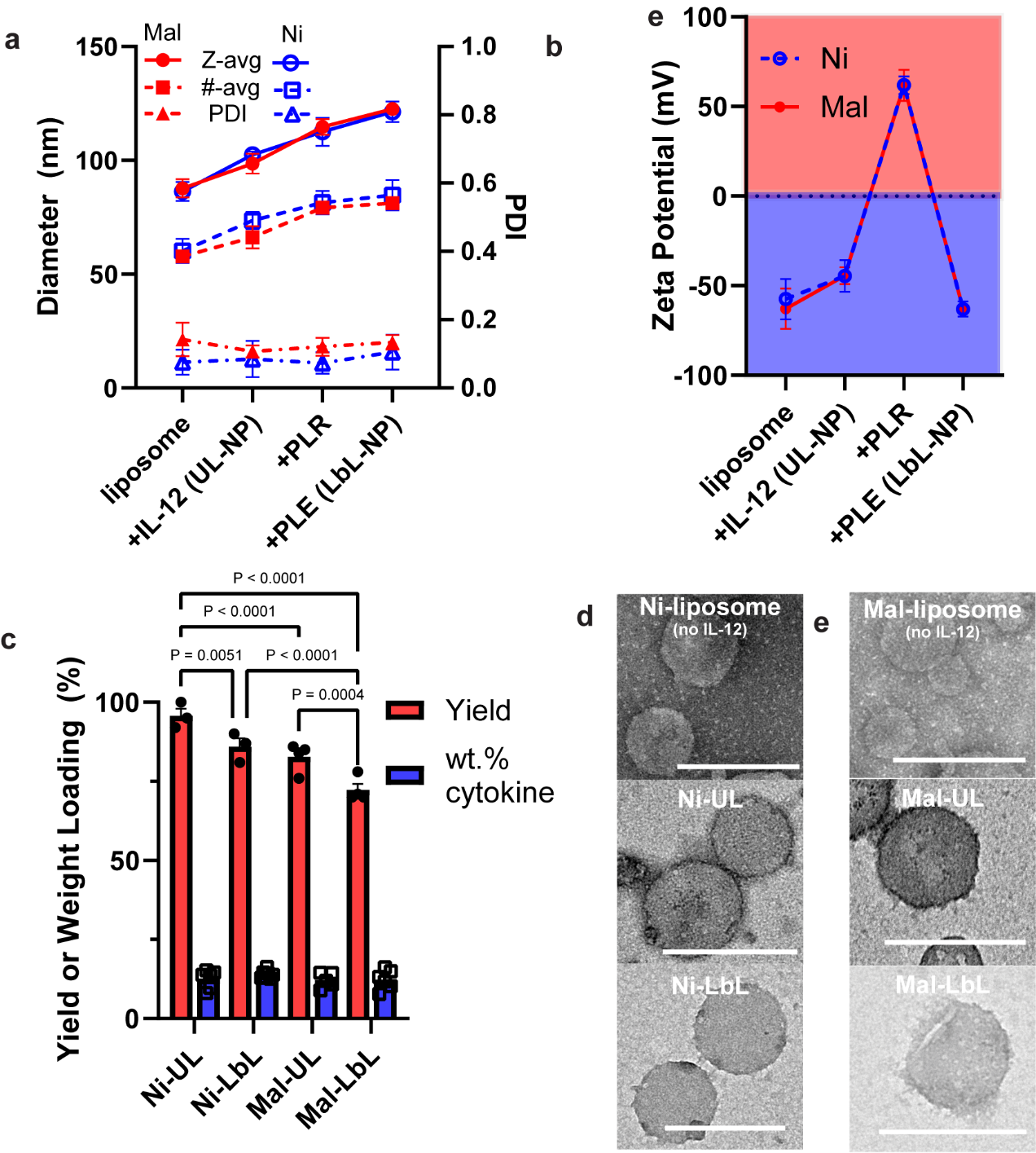

**Figure S2. Synthesis of LbL-NPs conjugated with IL-12 via either maleimide-cysteine reaction or nickel-histidine interaction yield similar particle biophysical properties. a**, Intensity-weighted hydrodynamic size (Z-avg), number average size (#-avg), and polydispersity index (PDI) of NPs during synthesis as measured via dynamic light scattering (mean ± s.d.). **b,** Zeta potential of NPs during synthesis as measured via electrophoretic mobility in deionized water (mean ± s.d.). **c,** Yield and weight loading of IL-12 for unlayered and layered particles with nickel-histidine linker (Ni-UL and Ni-LbL) and unlayered and layered particles with a maleimide-cysteine bond (Mal-UL and Mal-LbL) (mean ± s.e.m.). **d-e,** Negative-stain (NS) transmission electron microscopy (TEM) with phosphotungstic acid of particles during synthesis with nickel-containing lipids (**d**) and maleimide-containing lipids (**e**) - scale bars represent 200 nm. Unlayered (UL) NPs without IL-12 presented the typical low-contrast micrographs of liposomes. When conjugated with IL-12, however, a dark rim around the liposomes could be observed which became diffuse after LbL deposition, suggesting successful IL-12 conjugation and LbL coating. Data are presented as mean values ± error with *n* = 3 independent batches of NPs. Statistical comparisons in **c** were performed using two-way analysis of variance (ANOVA) with Tukey’s multiple-comparisons test.

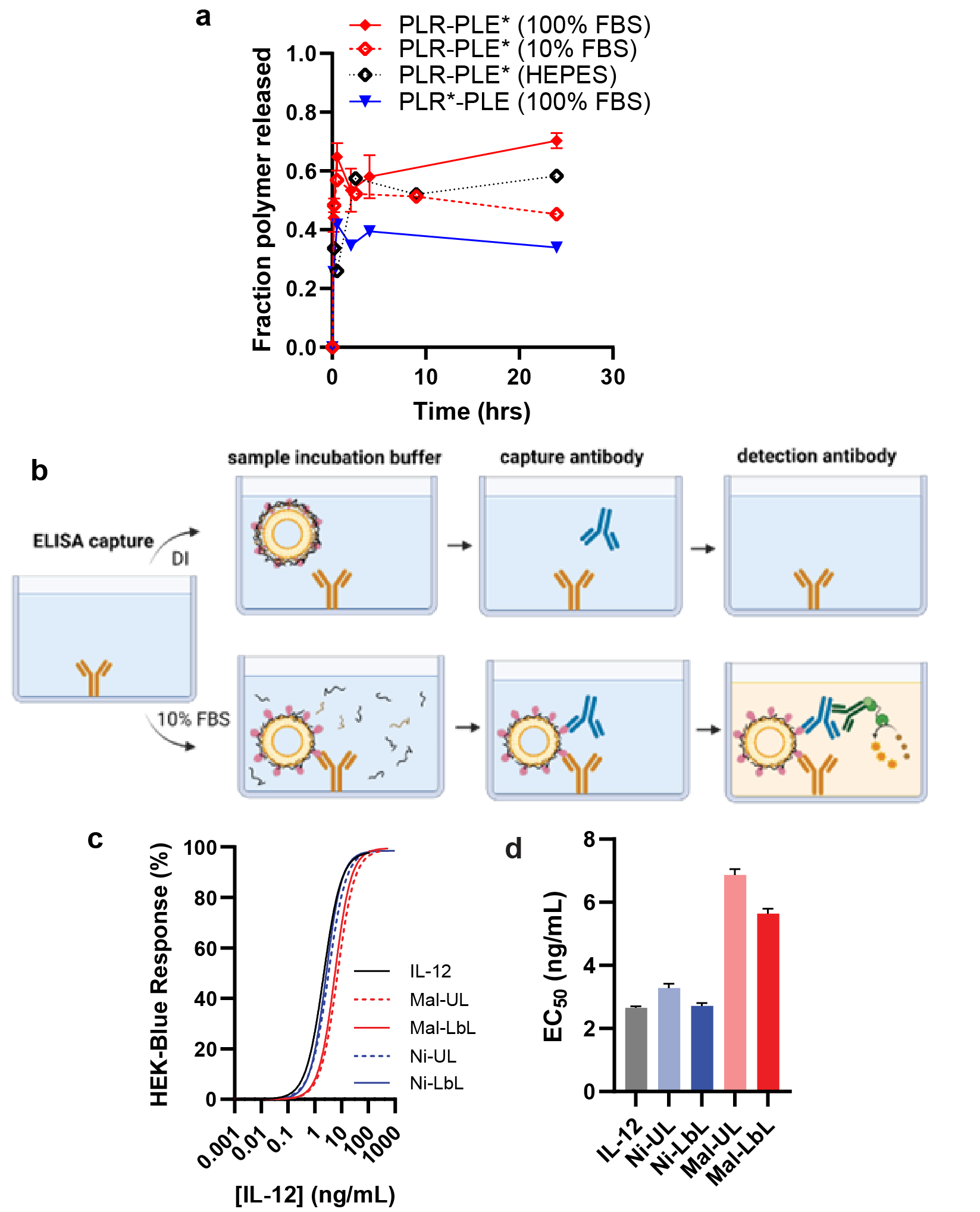

**Figure S3. Polyelectrolyte film is partially released in buffers with physiological ionic strength and does not block IL-12 availability in LbL-NPs. A,** Measurement of PLE or PLR release from LbL film on NPs incubated at 37 °C in 15 mM HEPES 150 mM NaCl (HEPES), 10%, or 100% FBS (mean ± s.e.m). **b,** Schematic for monoclonal antibody capture of NP-bound IL-12 and detection in varying buffer conditions. **c,** HEK-Blue IL-12 reporter cell line response to IL-12 in various formats (*n*>100 points per curve from 7 independent particle batches). (**d**) Calculated IL-12 EC_50_ from HEK-Blue IL12 response curves (mean ± s.e.m).

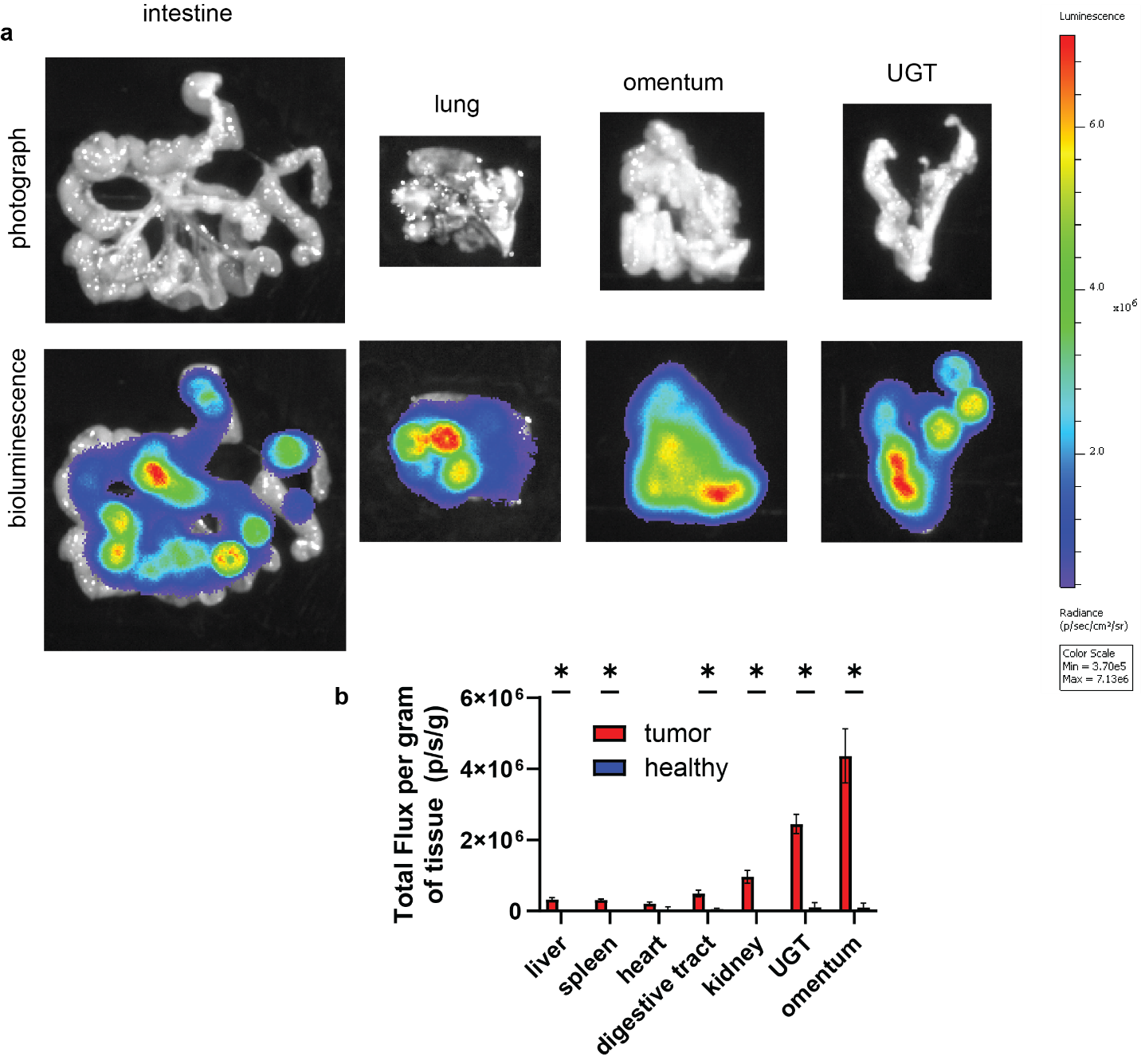

**Figure S4. Characterization of metastatic ovarian cancer model - OV2944-HM1. a-b,** B6C3F1 mice were inoculated with 10^6^ HM-1-luc tumor cells i.p. on day 0 (*n* = 40) or left as naïve healthy animals (*n* = 10). Shown are representative, intestine, lung, omentum and UGT ex-vivo IVIS BLI images (**a**) and quantitation of BLI signal in healthy organs compared to organs two weeks after tumor inoculation (mean ± s.e.m, **b**). Statistical comparisons in **b** were performed using the nonparametric Mean-Whitney test with correction for multiple comparisons based on a false discovery rate of 1% (q=0.0136).

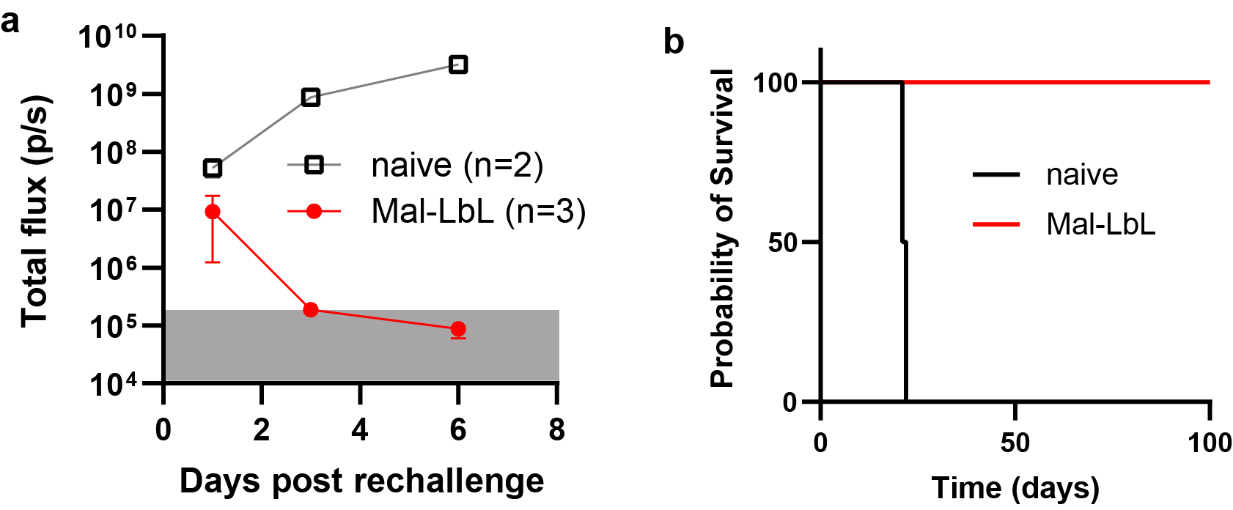

**Figure S5. Mice with complete remission of metastatic ovarian cancer demonstrate strong immune memory induction upon i.p. luc-HM-1 rechallenge. a-b**, B6C3F1 mice (*n* = 10/group) inoculated with 10^6^ HM-1-luc tumor cells on day 0 were treated on days 7 and 14 with 20 µg of IL-12 as a free cytokine or conjugated to NPs. On day 100, surviving Mal-LbL mine (*n* = 3) or naïve (*n* = 2) were injected with 3x10^5^ luc-HM-1 cells i.p. Shown are in vivo IVIS whole-animal i.p. BLI readings (mean ± s.e.m., **b**), and overall survival (**c**).

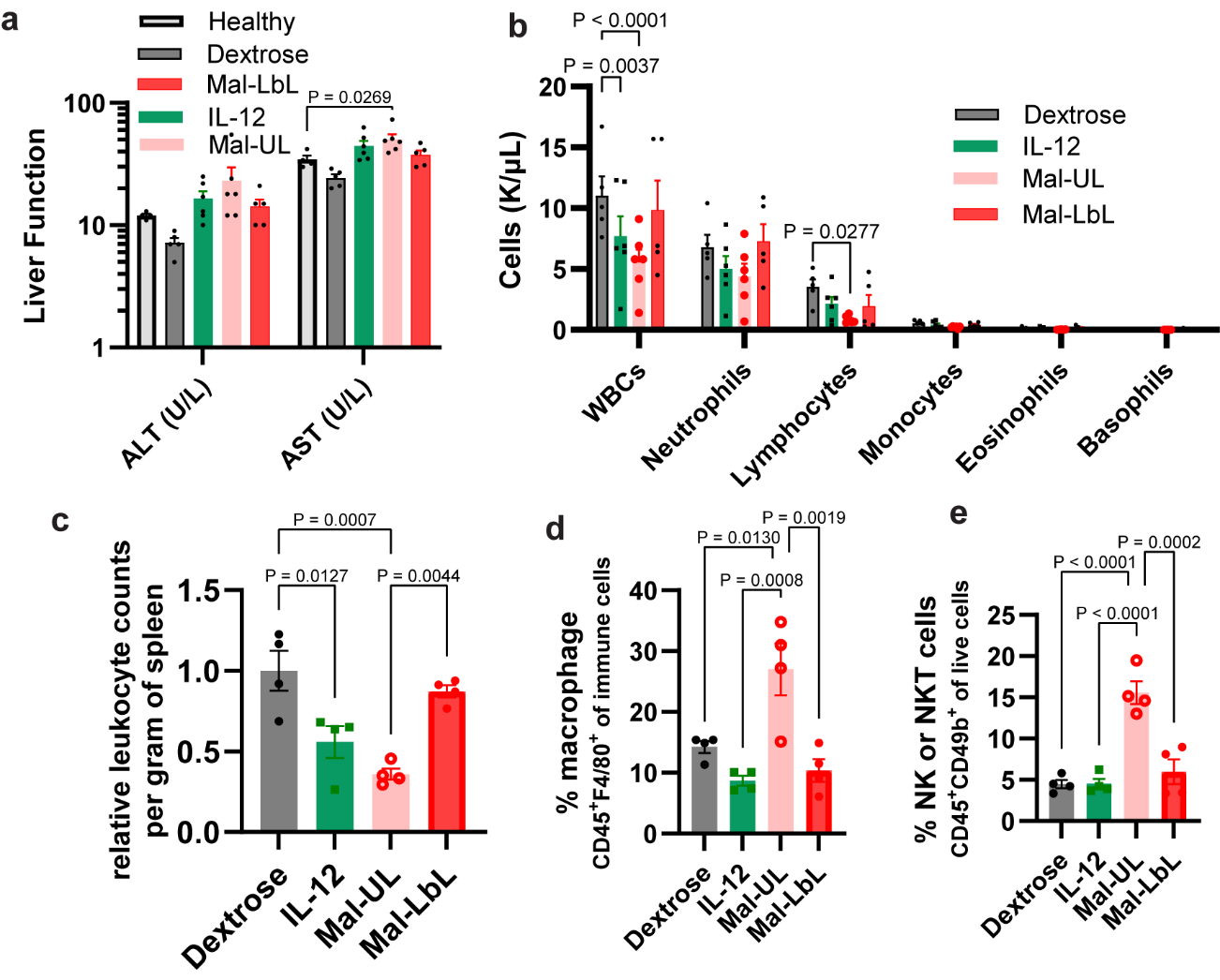

**Figure S6. Intraperitoneal dosing of 20 µg of Mal-LbL NPs does not cause systemic toxicity. a-e**, B6C3F1 mice inoculated with 10^6^ HM-1 tumor cells on day 0 were treated on days 10 with 20 µg of IL-12 as a free cytokine or conjugated to Mal NPs (UL and LbL). Two days after dosing blood (n = 6/group) and spleens (n = 4/group) were harvested and sent for a complete blood panel or processed for flow cytometry analysis, respectively. Shown are serum levels of liver damage markers (alanine transaminase – ALT - and aspartate aminotransferase - AST) compared to healthy mice controls (*n* = 4, **a**), complete blood count panel (**b**), quantitation of live leukocyte (CD45^+^) counts in spleen (**c**), and percentage of macrophage (**d**) and NK (**e**) cell in splenocytes. Statistical comparisons performed using two-way (**a,b**) or one-way (**c,d,e**) analysis of variance (ANOVA) with Tukey’s multiple-comparisons (liver enzyme measurement was compared to healthy controls).

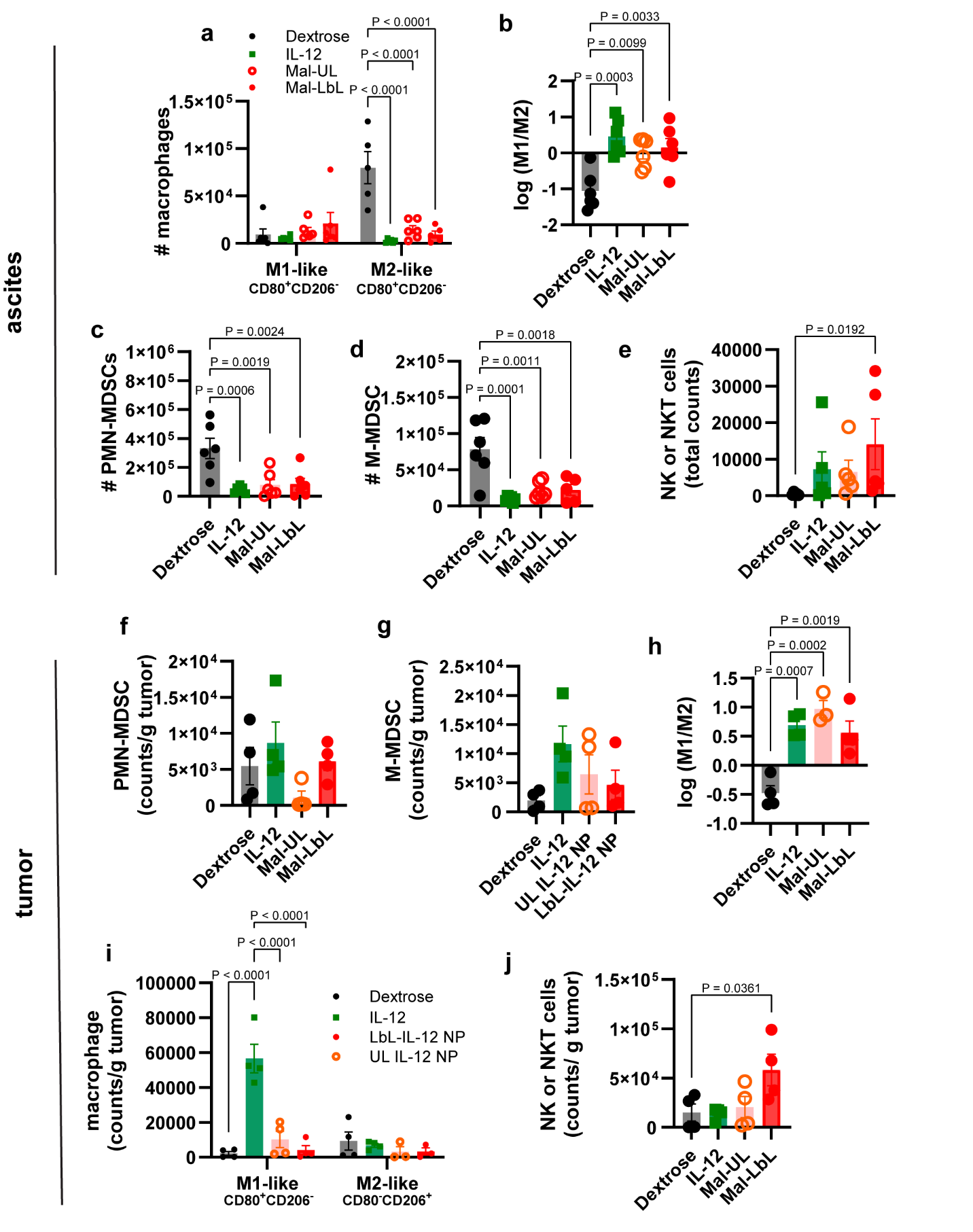

**Figure S7. Immune phenotyping of cells in ascites fluid and tumor tissues. a-j**, B6C3F1 mice inoculated with 10^6^ HM-1 tumor cells on day 0 were treated on days 10 with 20 µg of IL-12 as a free cytokine or conjugated to Mal NPs (UL and LbL). Two days after dosing ascites (*n* = 6/group) and i.p. tumor nodules (primarly omentum tissue, *n* = 4/group) were harvested and processed for flow cytometry analysis. Shown are total counts of M1-like and M2-like macrophages **(a)**, logarithmic of M1-like to M2-like macrophages ratio (**b**), and total counts of PMN-MDSC (**c**), M-MDSC (**d)**, and NK cells (**e**) in ascites fluid. Also shown are total counts of PMN-MDSC (**f**) and M-MDSC (**g**), logarithmic of M1-like to M2-like macrophages ratio (**h**), and total counts of M1-like and M2-like macrophages (**i**), and NK cells (**j**) in tumor nodules. Statistical comparisons were performed using two-way (**a,i**) or one-way (**b,c,d,e,h,j**) analysis of variance (ANOVA) with Tukey’s multiple comparisons (liver enzyme measurement was compared to healthy controls).

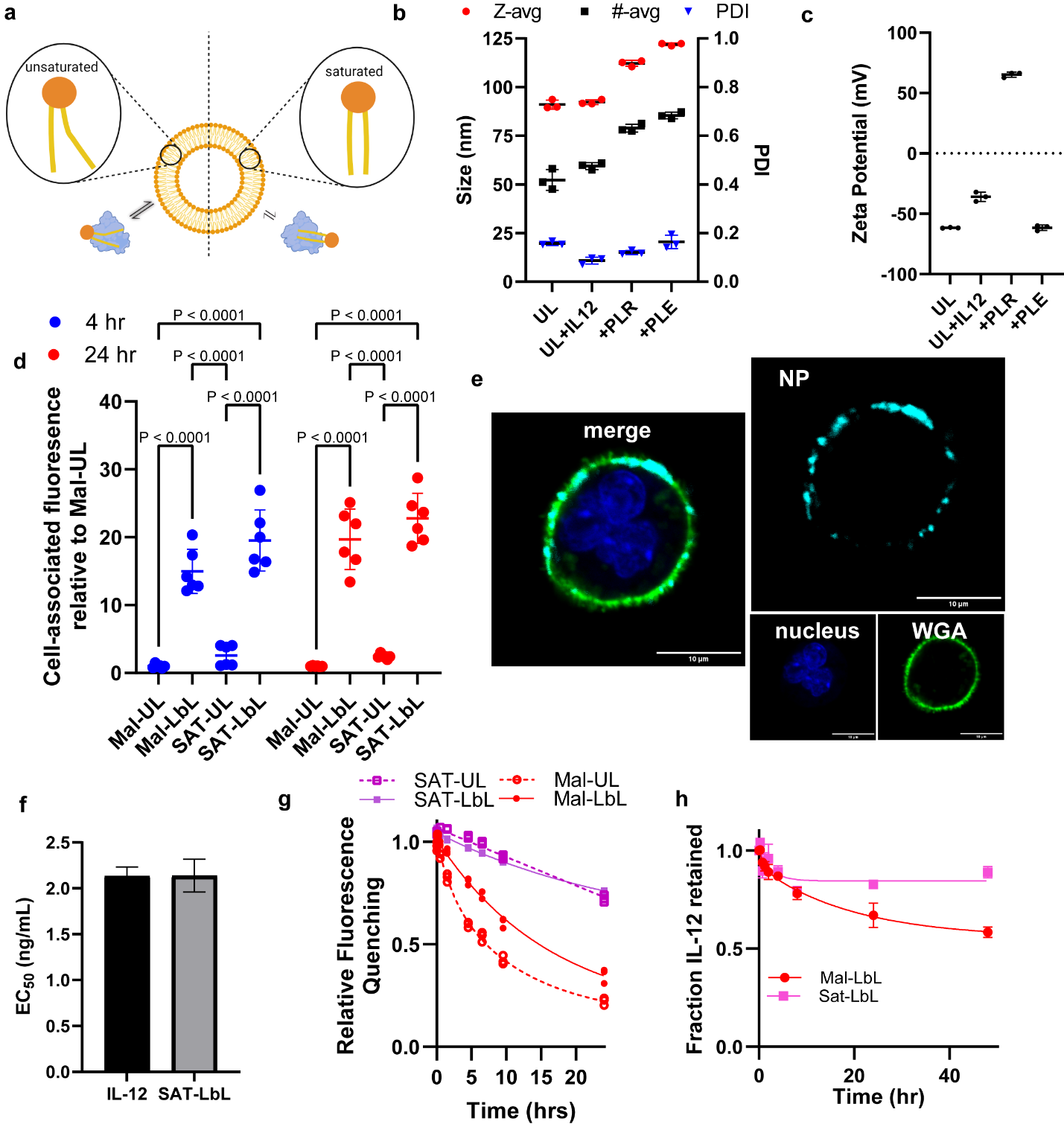

**Figure S8. Characterization of Mal IL-12 NPs composed of saturated (SAT) lipids. a**, Illustration of liposome bilayer composition effect on lipid exchange rate with serum proteins. **b,** Intensity-weighted hydrodynamic size (Z-avg), number average size (#-avg), and PDI of NPs during synthesis as measured via DLS (mean ± s.d.). **c,** Zeta potential of NPs during synthesis as measured via electrophoretic mobility in deionized water (mean ± s.d.). **d,** Association of NP fluorescence with HM1 cells *in vitro* relative to unlayered NPs after 4 and 24 hours of incubation (mean ± s.d.). **e,** Representative confocal microscopy images of HM-1 cells dosed with SAT-LbL for 4 hours. **f,** Calculated IL-12 EC_50_ of IL-12 compared to SAT-LbL NPs from HEK-Blue IL-12 assay (mean ± s.e.m.). **g,** Assessment of de-quenching from fluorophore detachment from unsat and SAT NPs when incubated with 100% FBS at 37 °C – curves represent the best fit of a two-phase decay model. **h,** Quantification of IL-12 retention with Mal-LbL or SAT-LbL upon incubation with 100% FBS (mean ± s.e.m.) – curves represent the best fit of a two-phase decay model. Statistical comparisons performed in **d** using two-way analysis of variance (ANOVA) with Tukey’s multiple-comparisons. Data are representative of at least two independent experiments.

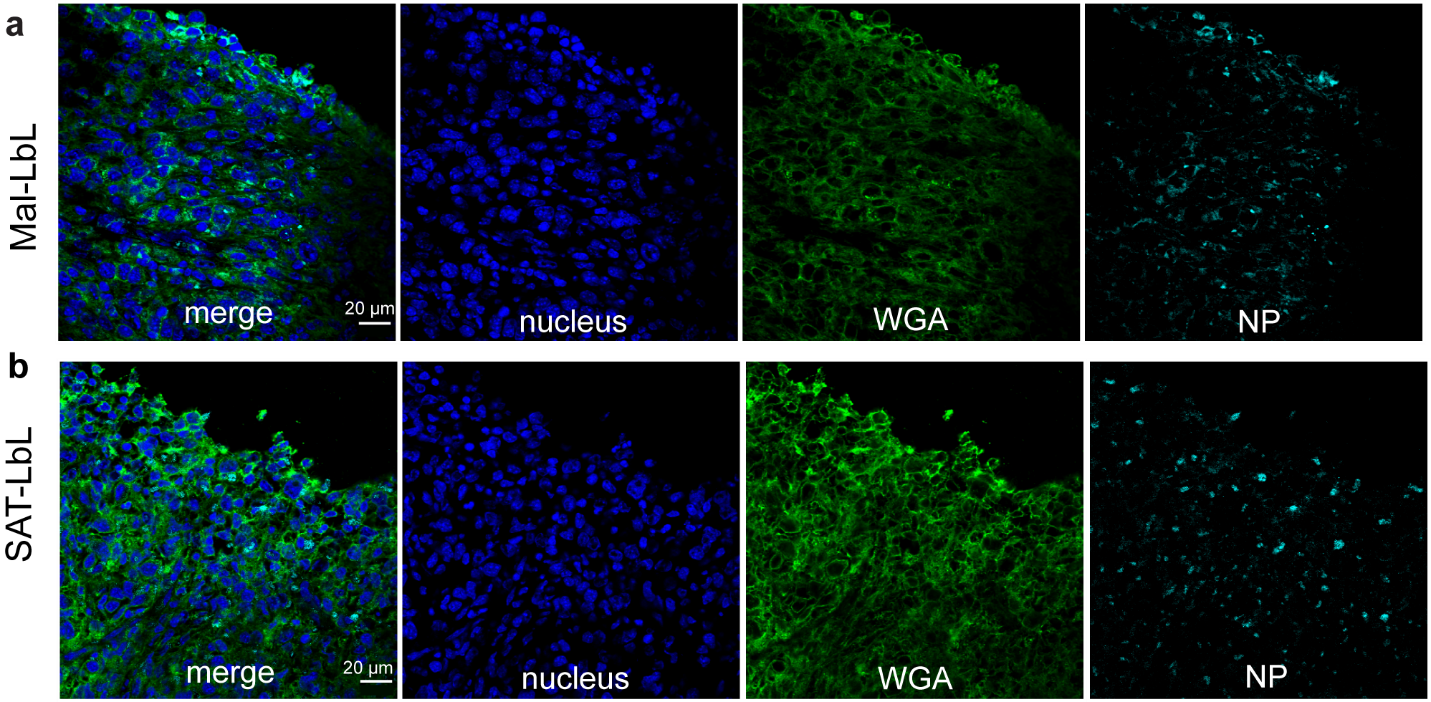

**Figure S9. Confocal microscopy analysis of histological cryosections of omentum tumor nodules demonstrates both Mal-LbL and SAT-LbL penetrate tumor tissue. a-b,** B6C3F1 mice were inoculated with 10^6^ HM-1-luc tumor cells on day 0 were administered fluorescently-tagged Mal-LbL of SAT-LbL NPs carrying 20 µg IL-12 on day 14. One day after dosing, animals were sacrificed, and the omentum containing tumor nodules was frozen in optimal cutting temperature (OCT) compound then frozen sectioned and stained for confocal microscopy analysis. Shown are representative confocal images of omental tumor nodules from Mal-LbL (**a**) and SAT-LbL (**b**) treated animals.

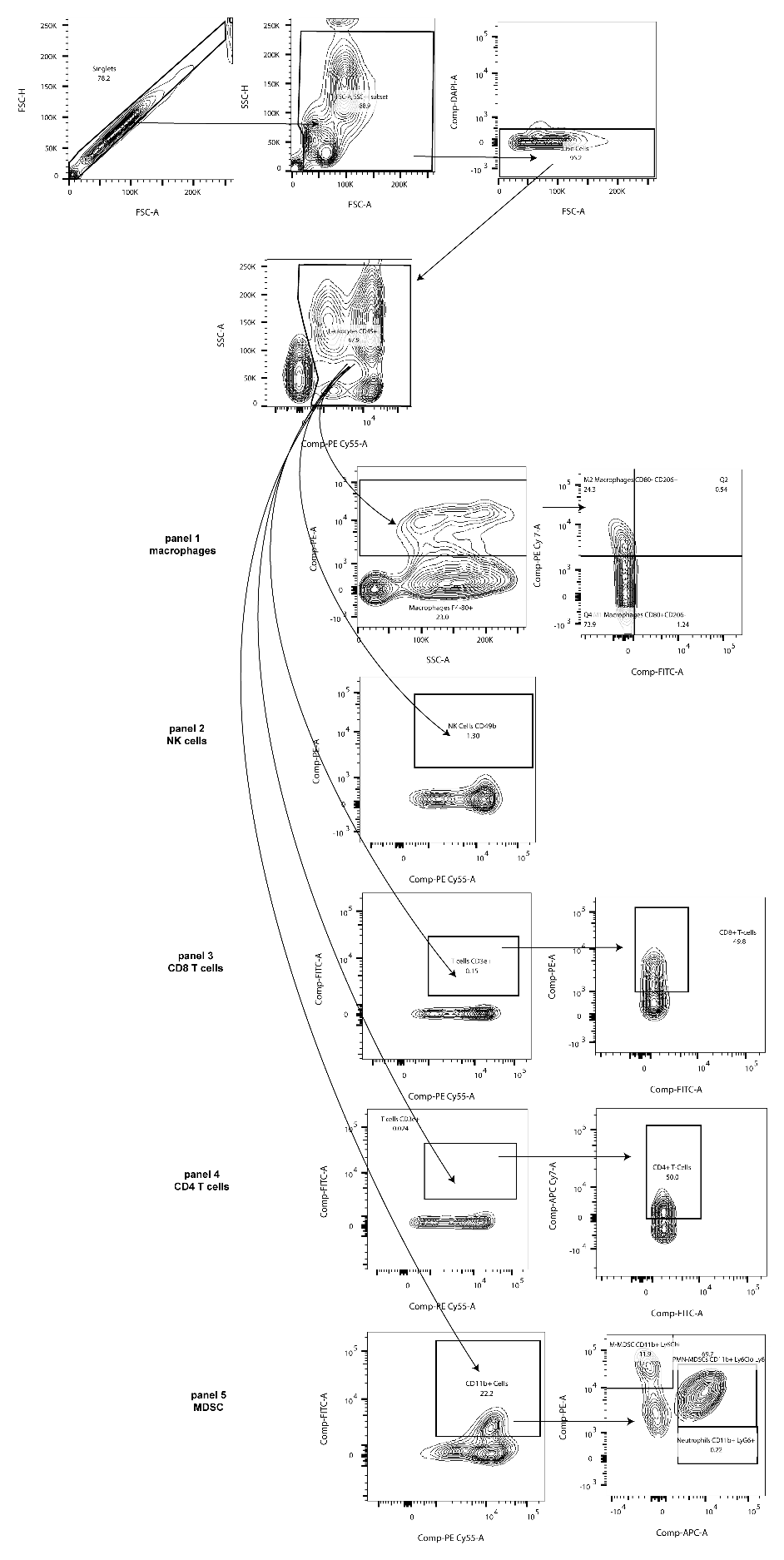

**Fig. S10.** Flow cytometry cell gating strategy.
